# EVALUATING METHODS AND PROTOCOLS OF FERRITIN-BASED MAGNETOGENETICS

**DOI:** 10.1101/2020.12.10.419911

**Authors:** Miriam Hernández-Morales, Victor Han, Richard H Kramer, Chunlei Liu

**Affiliations:** Department of Electrical Engineering and Computer Sciences, University of California, Berkeley, CA 94720, USA; Helen Wills Neuroscience Institute, University of California, Berkeley, CA 94720, USA; Department of Molecular and Cell Biology, University of California, Berkeley, Berkeley, CA 94720, USA

**Keywords:** Magnetogenetics, neuromodulation, radiofrequency magnetic fields

## Abstract

FeRIC (Ferritin iron Redistribution to Ion Channels) is a magnetogenetic technique that uses radio frequency (RF) alternating magnetic fields to activate the transient receptor potential channels, TRPV1 and TRPV4, coupled to cellular ferritins. In cells expressing ferritin-tagged TRPV, RF stimulation increases the cytosolic Ca^2+^ levels via a biochemical pathway. The interaction between RF and ferritin increases the free cytosolic iron levels that in turn, trigger chemical reactions producing reactive oxygen species and oxidized lipids that activate the ferritin-tagged TRPV. In this pathway, it is expected that experimental factors that disturb the ferritin expression, the ferritin iron load, the TRPV functional expression, or the cellular redox state will impact the efficiency of RF in activating ferritin-tagged TRPV. Here, we examined several experimental factors that either enhance or abolish the RF control of ferritin-tagged TRPV. The findings may help optimize and establish reproducible magnetogenetic protocols.

## Introduction

Magnetogenetics is a terminology loosely used to describe a group of techniques that apply magnetic fields to control cell activity via interactions with certain proteins. This group of techniques uses a diverse range of magnetic field conditions to target various choices of membrane proteins. These differences have contributed to the confusion and debates regarding the biophysical mechanisms and experimental reproducibility. In one class of magnetogenetic techniques, static, low frequency or radio frequency (RF) alternating magnetic fields have been applied to activate transient receptor potential vanilloid channels (TRPV) coupled to ferritin (Brier et al., 2020; Duret et al., 2019; Hernández-Morales et al., 2020; Hutson et al., 2017; Stanley et al., 2016). The biophysical mechanisms responsible for magnetic activation of the channels are still being worked out. It was first proposed that the interaction between static or RF magnetic fields and ferritin produces heat or mechanical stimuli that directly activate TRPV1 and TRPV4, which are intrinsically temperature- and mechanically-sensitive (Stanley et al., 2016; Wheeler et al., 2016). However, no change in temperature could be detected at the surface of ferritin upon RF exposure (Davis et al., 2020). Likewise, a theoretical calculation estimates that the temperature change produced by interaction of magnetic fields with ferritin is several orders of magnitude lower than that required to activate TRPV (Meister, 2016). Other theoretical estimations, assuming the ferritin is at the maximum iron load (∼ 4500 iron atoms) and has specific magnetic properties (superparamagnetic), have proposed potential mechanisms, such as the magnetocaloric effect, that might contribute to magnetic heating and subsequent activation of TRPV (Barbic, 2019; Duret et al., 2019). However, there is no conclusive experimental evidence supporting those mechanistic proposals. Recently, we proposed an indirect biochemical mechanism that allows RF to activate ferritin-tagged TRPV. Specifically, by triggering dissociation of iron from the ferritin, RF catalyzes the generation of reactive oxygen species (ROS), short-chain fatty acids, and oxidized lipids, which are activators of TRPV (Hernández-Morales et al., 2020). This biochemical pathway and the role of ROS in the RF-induced activation of ferritin-tagged TRPV have been recently corroborated (Brier et al., 2020).

Besides the uncertainties about the underlying mechanisms, the efficiency of a specific magnetogenetic technique that uses a ferritin-fused TRPV4, named Magneto2.0 (Wheeler et al., 2016) has been questioned. Three independent groups reported the failure to activate neurons expressing Magneto2.0 upon stimulation with static magnetic fields (Kole et al., 2019; Wang et al., 2019; Xu et al., 2019). It has been proposed that diverse experimental factors are responsible for those discrepancies such as the magnetic stimuli (static versus low frequency magnetic fields), the viral vectors used to deliver Magneto2.0 (pAAV versus Semliki Forest virus, Sindbis virus, and lentivirus), and the transduction period to achieve functional expression of Magneto2.0 (Wheeler et al., 2020). However, there is no experimental evidence reported to support the hypothesis that those specific experimental factors are responsible for conflicting results using Magneto2.0.

The lack of a unified experimental protocol has compounded the many unresolved issues. For example, ferritin-based magnetogenetic approaches use diverse magnetic stimuli (static, low frequency, kHz RF, or MHz RF magnetic fields) and have tagged TRPV with both endogenous or chimeric ferritin (Brier et al., 2020; Duret et al., 2019; Hernández-Morales et al., 2020; Hutson et al., 2017; Stanley et al., 2016; Wheeler et al., 2016). Other factors that may contribute to the reported inconsistencies are the function and expression of both ferritin and TRPV. Ideally, ferritins should be at the maximum iron load to transduce magnetic fields proficiently. Nevertheless, it is unknown if the chimeric ferritins store iron at the same level as endogenous ferritins. Furthermore, the iron load of ferritins is highly variable from almost empty up to maximum load (∼4500 iron atoms) (Jian et al., 2016). Regarding TRPV, those channels should be functionally expressed at the cell membrane. However, their expression and sensitivity to diverse stimuli are subjected to cellular regulatory mechanisms. Several TRPV channels are constitutively active, and cells prevent the associated cytotoxic effect by downregulating the TRPV density at the plasma membrane (Ferrandiz-Huertas et al., 2014; Montell, 2004; Planells-Cases and Ferrer-Montiel, 2007).

Given the many potential variables described above, we reasoned that experimental factors that disturb the cellular iron homeostasis and the ferritin and TRPV channel expression may impact the magnetic control of ferritin-tagged TRPV. Using a single magnetogenetic approach, FeRIC technology, we examined the influence of diverse experimental variables on the RF control of cytosolic Ca^2+^ levels in cells expressing ferritin-tagged TRPV4. Interestingly, we found that while some experimental variables abolished magnetic control of ferritin-tagged TRPV4, others enhanced it. The observations reported here may contribute to the standardization and optimization of current and future magnetogenetic techniques to achieve better reproducibility.

## Results

### I. FeRIC technology allows the magnetic control of cytosolic Ca^2+^ levels

Here we examined the influence of different experimental factors on the Ca^2+^ responses induced by RF fields in cultured cells expressing the ferritin-tagged channel TRPV4^FeRIC^. RF fields at 180 MHz and 1.6 µT were generated with solenoid coils enclosing a cell culture dish (Figure S1A, B). The cytosolic Ca^2+^ levels were monitored in Neuro2a (N2a) cells expressing GCaMP6 plus TRPV4^FeRIC^. All experiments were performed at room temperature (22°C) except for those testing TRPV4^FeRIC^ responsiveness at 32°C and 37°C. The cells expressing TRPV4^FeRIC^ were identified using the mCherry reporter. Data were quantified as the change in GCaMP6 fluorescence divided by baseline fluorescence (ΔF/F0), the GCaMP6 area under the curve (AUC), and the fraction of cells responsive to RF (RF responsiveness). A cell was considered RF responsive when the GCaMP6 ΔF/F0 increased 10 times over the standard deviation of its baseline fluorescence (see **STAR methods**).

First, we estimated the distribution of the electric (E) and magnetic (B) fields applied to the cells to rule out the potential contribution of the E field to RF-induced Ca^2+^ responses. To estimate the distribution of the E and B fields, we computed them using the finite-difference time-domain (FDTD) method implemented by the openEMS project (Liebig et al., 2013). The simulation setup included the 5 cm-diameter RF coil containing the 3.5 cm-diameter dish half-filled with imaging saline solution (dish height: 1 cm, saline solution height: 0.5 cm) (Figure S1C). The estimated E field corresponding to a 180 MHz and 1.6 µT magnetic field, at the center of the culture dish, was about 5.5 V/m. The magnitude distributions in two dimensions, for both transverse and longitudinal cross-sections, of the B and E fields are shown in Figure 1A. As expected, the strength of the B field remains relatively large at the center of the coil/dish. In contrast, the E field amplitude drastically decreases in the center of the coil/dish. Although the estimated electric field values for the RF at 180 MHz and 1.6 µT are similar in amplitude to those needed for transcranial magnetic stimulation (TMS) (Fox et al., 2004; Zmeykina et al., 2020), they should be negligible and should not produce significant effects on the membrane ion channels. Firstly, the induced potential difference between the two opposite sides of a cell membrane-embedded ion channel exposed to an E = 5.5 V/m is about 28 nV (Vind= E*d*; *d* is membrane thickness = 5 nm). This change in the membrane potential should be insufficient to activate voltage-gated ion channels and, in addition, TRPV is only weakly voltage-dependent (Nilius et al., 2005). TMS-induced E-fields have been measured to be on the order of 200 V/m, which is still considered to underestimate the peak E-field amplitude due to the short duration of TMS pulses (Nieminen et al., 2015). While some TMS models predict channel activation through more complicated mechanisms despite E fields of only tens of V/m (Pashut et al., 2011), the frequency of a 180 MHz RF stimulus is four orders of magnitude higher compared to the kHz range of TMS. The effects of E fields on cell membrane capacitance are different at kHz and MHz frequencies. For N2a cells, a membrane capacitance of 10 pF (Gutiérrez-Martín et al., 2011) provides a reactance of about 1.6E5 ohms at 100 kHz, but at 180 MHz it is only about 88 ohms. With this greatly decreased impedance, the E fields associated with a 180 MHz RF likely produce negligible charge and voltage buildup on the cell membrane.

**Figure 1.**
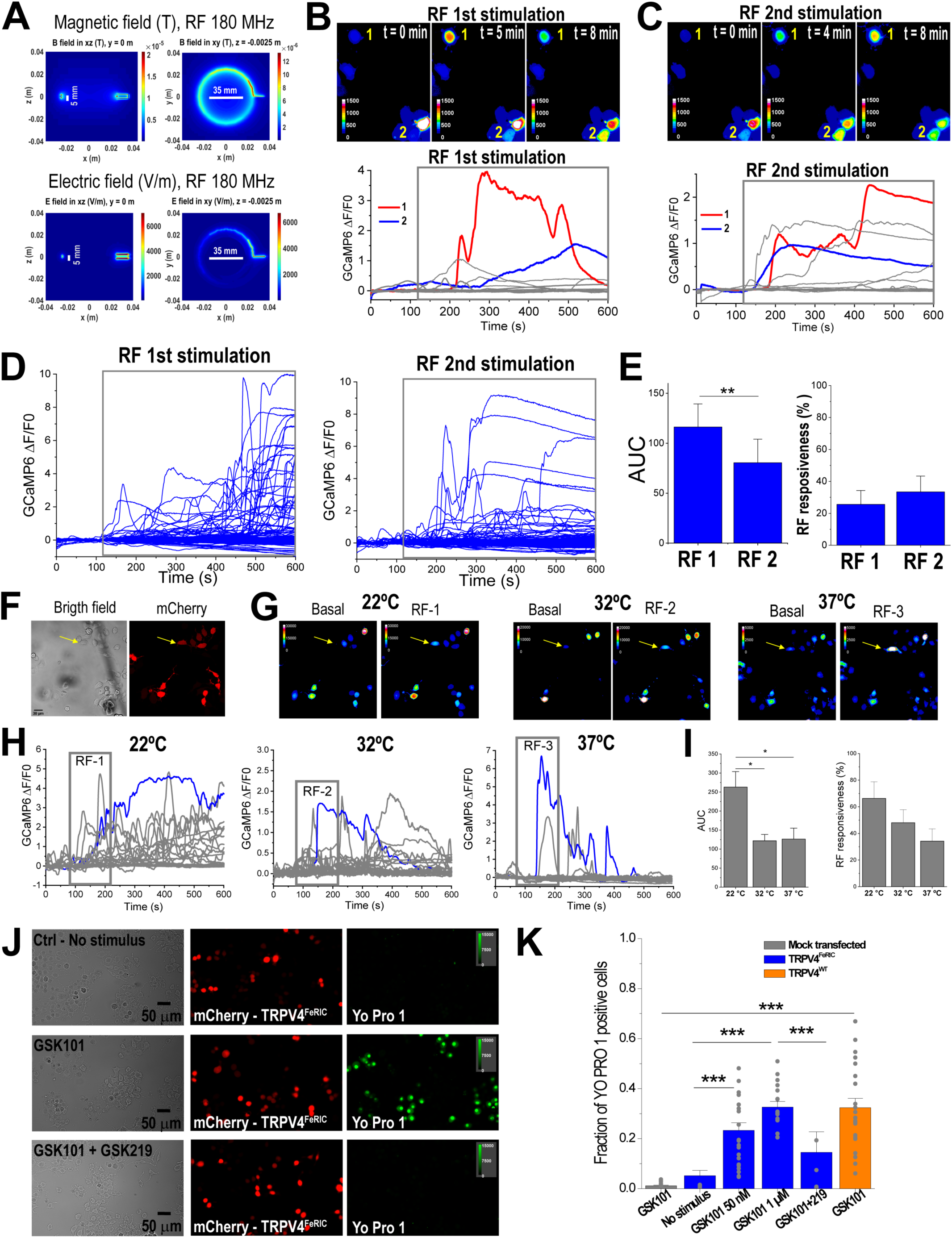
Consecutive RF stimulation at MHz frequency activates TRPV4^FeRIC^ at different temperatures. (**A**) Distribution of the magnetic and electric fields produced by RF fields (180 MHz, 1.6 µT) in a culture dish. Vertical lines: 5 mm height of the saline solution. Horizontal lines: 35 mm-diameter of the culture dish. Cultured cells are at the center of the culture dish. (**B** - **C**) Top images are pseudocolor images of GCaMP6 fluorescence and bottom plots are the changes in GCaMP6 ΔF/F0 of N2a cells expressing TRPV4^FeRIC^ before and upon the (**B**) 1^st^ and (**C**) 2^nd^ RF stimulations. Cells 1 and 2 were responsive to 1^st^ and 2^nd^ RF stimulations. (**D**) Changes in GCaMP6 ΔF/F0 from all tested N2a cells expressing TRPV4^FeRIC^ stimulated with two consecutive RF stimulations (gray boxes). (**E**) Average changes (± SEM) in GCaMP6 AUC and RF responsiveness for the two consecutive RF stimulations. (**F**) Bright-field and fluorescence images of N2a cells expressing TRPV4^FeRIC^ (mCherry+). Scale bar = 20 µm. (**G**) Pseudocolor images of GCaMP6 fluorescence from cells in (F), before and after RF stimulation at 22°C, 32°C, and 37°C. (**H**) Changes in GCaMP6 ΔF/F0 and (**I**) average changes (± SEM) in GCaMP6 AUC and RF responsiveness from N2a cells expressing TRPV4^FeRIC^ stimulated consecutively with RF (2 min, gray boxes) at 22°C, 32°C, and 37°C. A cell that was activated with RF at all tested temperatures is indicated with yellow arrows in images and with blue traces in the plots. (**J**) Images of N2a cells expressing TRPV4^FeRIC^ (mCherry+) without stimulation or following GSK101 (50 nM) application for 20 min in the absence or the presence of GSK219. Scale bars = 50 µm. (**K**) Averaged fraction of N2a cells expressing TRPV4^FeRIC^ or TRPV4^WT^ that become permeable to Yo Pro 1 under different experimental conditions. For this and the following figures, significance was determined using one-way ANOVA followed by Tukey’s multiple comparisons test. Where applicable, either p<0.05 (*), p<0.001 (**), or p<0.0001 (***) was considered a statistically significant difference. See also Figures S1, S2 and Table S1.

It has been reported that RF fields at high kHz frequencies activate a ferritin-tagged TRPV1 (Brier et al., 2020; Stanley et al., 2015, 2016). Thus, we also estimated the E field amplitude for a 465 kHz RF stimulus of 31 μT which resulted in about 0.1 V/m (Figure S2A, B). As expected from Maxwell’s equations, the E field estimated for the kHz RF is smaller than the E field for the MHz RF, so it should also produce negligible effects on the cell membrane. We did not use a 465 kHz RF stimulus at a larger strength because starting at ∼200 μT it produces a focus drift during imaging experiments. This is consistent with a heating effect produced for a ∼500 kHz RF at mT strength (Brier et al., 2020). In contrast, the RF at 180 MHz and 1.6 μT stimulus did not increase the temperature in the saline imaging solution when applied for up to 20 min (Temperature initial: 22.09°C; Temperature final: 22.03°C; ΔT: -0.06°C).

Next, we evaluated the ability of FeRIC technology to consistently activate the ferritin-tagged TRPV4^FeRIC^ with consecutive RF stimulations even at different temperatures. As previously reported, RF stimulation (180 MHz at 1.6 μT for 8 min) increased the cytosolic Ca^2+^ levels in N2a cells expressing TRPV4^FeRIC^ (Figure 1B). After a 20-min recovery period, a second RF stimulation increased the cytosolic Ca^2+^ levels in those N2a cells (Figure 1C). Most of the TRPV4^FeRIC^-expressing N2a cells that were activated by the first RF stimulation were responsive to the second one (Figure 1D). The RF responsiveness between the 1st and 2nd RF stimulation was similar (1^st^: 25.6 ± 8 %; 2^nd^ RF: 33.4 ± 8 %) (Figure 1E); but the GCaMP6 AUC induced with the second RF stimulation was smaller with respect to the first one (1^st^ RF: 116.3 ± 23, n=151 cells; 2^nd^ RF: 80.5 ± 23.5, 6 experiments, 209 cells; p<0.001) (Figure 1E). In contrast, RF at 465 kHz and 31 μT only slightly increased the GCaMP6 fluorescence in TRPV4^FeRIC^-expressing N2a cells relative to unstimulated cells (Figure S2C; No RF 24-h: -17.4 ± 6.1; RF 24-h: 25.6 ± 8.6; p<0.05). The failure to activate Ca^2+^ responses with RF at 465 kHz and 31 μT may be explained because the strength (B) and frequency (f) product of this stimulus is ∼20 times lower compared with the B*f product of RF at 180 MHz and 1.6 μT. Several magnetogenetics studies have used RF at kHz frequencies, but in our experimental conditions, MHz RF showed clear advantages in terms of RF power, ease of setup, absence of RF-induced heating effect, and consistent activation of cells expressing TRPV4^FeRIC^, so we focused on MHz RF for the majority of this paper.

Next, because TRPV4 is activated at about 34°C (Caterina et al., 1997) and even at temperatures as low as 32°C (Chung et al., 2003), we examined the RF-induced activation of TRPV4^FeRIC^ at 32°C and 37°C. In a series of experiments, N2a cells expressing TRPV4^FeRIC^ were consecutively stimulated with RF (180 MHz, 1.6 μT) at 22°C, 32°C, and 37°C (Figure 1F-H). Because increasing the temperature from 22°C to 32-37°C activates TRPV4^FeRIC^, cells were rested for ∼20 min before applying the RF stimulus. In N2a cells expressing TRPV4^FeRIC^, RF stimulation increased the cytosolic Ca^2+^ levels at all tested temperatures (Figure 1G, I). Although the RF responsiveness was smaller at 32°C and 37°C compared to 22°C, the difference was not statistically significant (22°C: 66.3 ± 12.5 %; 32°C: 48 ± 9.5; 37°C: 34.2 ± 9.4; n = 56 cells, 4 independent experiments) (Figure 1I). Interestingly, RF-induced Ca^2+^ responses at 37°C displayed faster activation and decay kinetics relative to those at either 22°C or 32°C. The faster kinetics of Ca^2+^ responses produced a significant decrease in the GCaMP6 AUC (22°C: 263 ± 40.8; 32°C: 121.9 ± 16.6; 37°C: 126.2 ± 28.8; n = 4 independent experiments, 56 cells) (Figure 1H, I).

To corroborate the expression of FeRIC channels at the cell membrane, we examined the permeability of N2a cells expressing TRPV4^FeRIC^ to Yo Pro 1, which is a large cation that binds to DNA. TRPV are cation channels permeable to Na^+^, K^+^, and Ca^2+^; however, after continuous agonist stimulation, some members of the TRPV family become permeable to larger cations, such as Yo Pro 1, due to a pore dilation process (Banke et al., 2010; Ferreira and Faria, 2016; McCoy et al., 2017). Stimulation with the TRPV4 agonist GSK 1016790A (GSK101) at 50 nM for 20 min of TRPV4^FeRIC^- and TRPV4^WT^-expressing N2a cells (at 24 h post-transfection) produced Yo Pro 1 influx in about 20 - 30% of examined cells (Figure 1J, K; TRPV4^FeRIC^: 21.3 ± 2.9 %, n = 3723 cell, 4 independent experiments; TRPV4^WT^: 32.4 ± 3.7 %, n = 3470 cells, 5 independent experiments). In contrast, Yo Pro 1 influx was not observed in mock-transfected N2a cells (1.3 ± 0.2 %, n =3001 cells, 4 independent experiments) or in TRPV4^FeRIC^-expressing N2a cells in the absence of GSK101 (no stimulus: 0.3 ± 0.1 %, n = 3448 cells, 4 independent experiments) or in the presence of the TRPV4 antagonist GSK 2193874 (GSK219; 3.1 ± 2.2 %, n = 1388 cells, 3 independent experiments) (Figure 1J, K).

These results indicate that a 180 MHz RF at 1.6 µT stimulus likely produces negligible electrical effects on membrane ion channels but consistently activates ferritin-tagged TRPV4 at physiologically relevant temperatures.

### II. RF-induced Ca^2+^ responses in TRPV4^FeRIC^-expressing N2a cells decreases with longer periods of TRPV4^FeRIC^ transient expression

In the literature, three main groups have reported successful magnetic control of ferritin-tagged TRPV channels expressed in diverse cultured cells (Brier et al., 2020; Hernández-Morales et al., 2020; Hutson et al., 2017; Stanley et al., 2016; Wheeler et al., 2016). However, the *in vitro* protocols used among those studies vary in time delays between seeding, transfection, and Ca^2+^ imaging. We examined the efficiency of RF stimulation (180 MHz at 1.6 μT) in inducing Ca^2+^ responses in TRPV4^FeRIC^-expressing N2a cells using adaptations of those three protocols. Timing protocol 1 (cells imaged 24-h post-transfection) is that we previously reported using FeRIC channels (Hernández-Morales et al., 2020; Hutson et al., 2017). Timing protocol 2 (cells transfected in a flask following plating for imaging) was adapted from the Magneto2.0 report (Wheeler et al., 2016). Timing protocol 3 (cells imaged 72-h post-transfection) was adapted from the anti-GFP–TRPV1/GFP–ferritin report (Stanley et al., 2016) (Figure 2A). In N2a cells imaged 24-h after transfection with TRPV4^FeRIC^ (timing protocol 1), RF stimulation increased the GCaMP6 AUC with respect to unstimulated cells (Figure 2B, C; Table S1; No RF: -17.4 ± 6.2; RF: 226.4 ± 31.4; p<0.0001). In contrast, in TRPV4^FeRIC^-expressing N2a cells imaged 48 or 72-h post-transfection (timing protocols 2 and 3, respectively), RF stimulation did not significantly increase the GCaMP6 fluorescence relative to unstimulated cells (Figure 2B, C; Table S1; No RF 48-h: 8.5 ± 12.8; RF 48-h: 95.5 ± 34.1; No RF 72-h: 21.1 ± 5.8; RF 72-h: 48.2 ± 26.2). For all protocols, RF stimulation did not change the cytosolic Ca^2+^ levels in N2a cells expressing only GCaMP6 and the functional expression of TRPV4^FeRIC^ was corroborated with GSK101 at 1 μM. Moreover, the RF-induced activation of TRPV4^FeRIC^ was inhibited with 1 μM GSK219 (Figure 2C).

**Figure 2.**
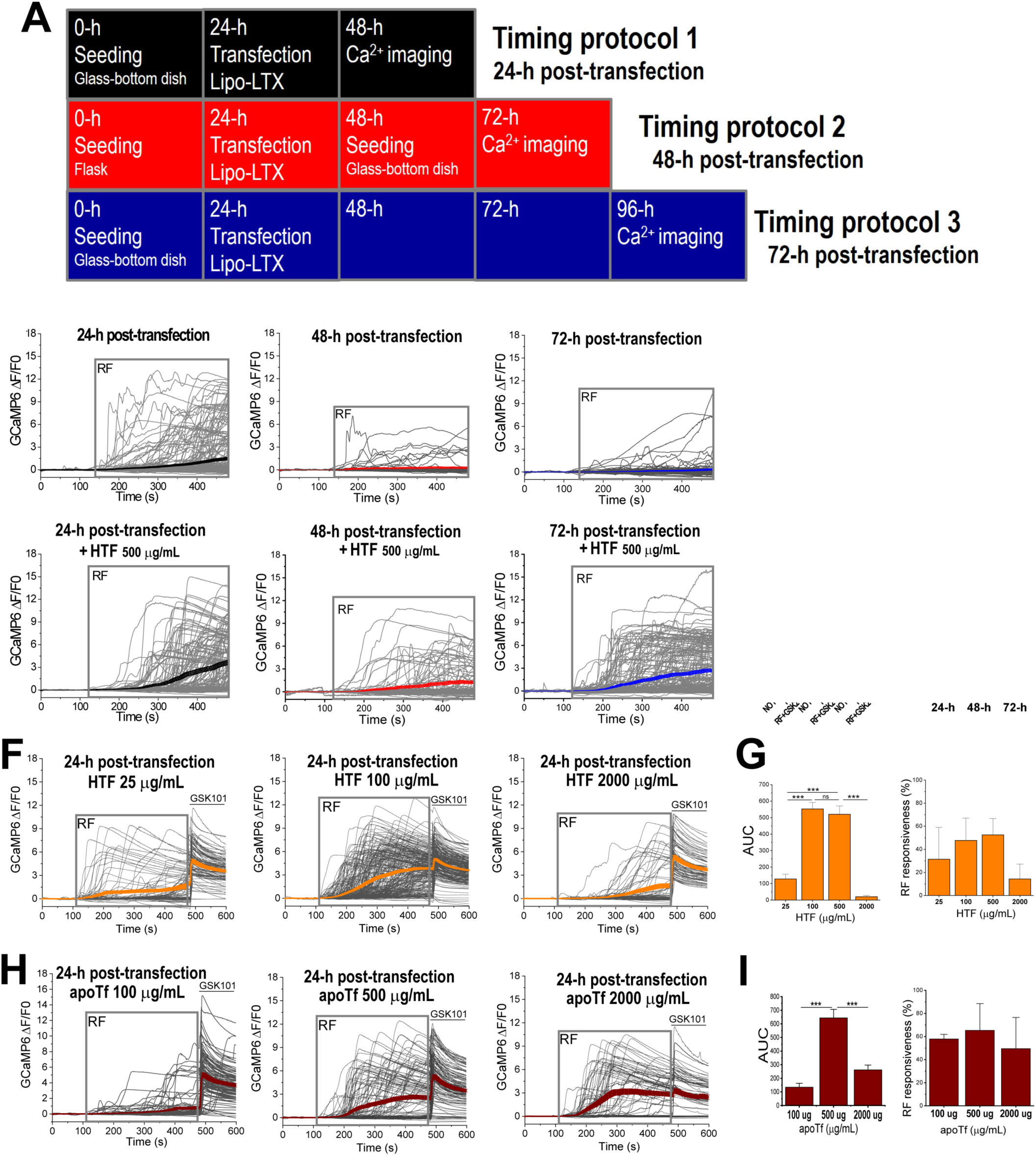
The period of expression of TRPV4^FeRIC^ and the cellular iron import influence the RF activation efficiency of TRPV4^FeRIC^. (**A**) Illustration of timing protocols used for testing RF-induced Ca^2+^ responses in N2a cells expressing TRPV4^FeRIC^. (**B**) Changes in GCaMP6 ΔF/F0 from TRPV4^FeRIC^-expressing N2a cells following RF stimulation (gray box). The average GCaMP6 ΔF/F0 (± SEM) is highlighted for cells imaged 24-h (black), 48-h (red), or 72-h (blue) post-transfection. (**C**) Average changes (± SEM) in GCaMP6 AUC and RF responsiveness from experiments in (B). (**D**) Changes in GCaMP6 ΔF/F0 from TRPV4^FeRIC^-expressing N2a cells treated with 500 μg/mL holotransferrin (HTF) following RF stimulation (gray box). The average GCaMP6 ΔF/F0 (± SEM) is highlighted for cells imaged 24-h (black), 48-h (red), or 72-h (blue) post-transfection. (**E**) Average changes (± SEM) in GCaMP6 AUC and RF responsiveness from experiments in (D). (**F**) Changes in GCaMP6 ΔF/F0 from TRPV4^FeRIC^-expressing N2a cells imaged 24-h after transfection and treated with 25, 100 or 2000 μg/mL HTF followed by RF stimulation (gray box). The average GCaMP6 ΔF/F0 (± SEM) is highlighted in orange. (**G**) Average changes (± SEM) in GCaMP6 AUC and RF responsiveness from experiments in (F). (**H**) Changes in GCaMP6 ΔF/F0 from TRPV4^FeRIC^-expressing N2a cells imaged 24-h after transfection and treated with 100, 500 or 2000 μg/mL apotransferrin (apoTF) followed by RF stimulation (gray box). The average GCaMP6 ΔF/F0 (± SEM) is highlighted in dark red. (**I**) Average changes (± SEM) in GCaMP6 AUC and RF responsiveness from experiments in (H). Where applicable, either p<0.05 (*), p<0.001 (**), or p<0.0001 (***) was considered a statistically significant difference. See also Figures S2, S3 and Table S1.

These results indicate that longer periods of TRPV4^FeRIC^ expression in N2a cells decrease their responsiveness to RF. This effect could be due to 1) a progressive decrease in the efficacy of ferritin as a RF transducer, 2) the functional downregulation of TRPV4^FeRIC^, 3) disturbance in the coupling between ferritin and TRPV4^FeRIC^, or 4) the combination of all these effects. Although it is unclear what is the cause of the loss of RF responsiveness over time, it is essential to determine the optimal time for expression of these channels when implementing magnetogenetic techniques with transient transfection.

### III. Increasing cellular iron import enhances the RF-induced Ca^2+^ responses in cells expressing TRPV4^FeRIC^

The efficiency of ferritin as the RF transducer depends on its iron load. To test if the ferritin iron load is involved in the loss of RF responsiveness with longer periods of TRPV4^FeRIC^ expression, N2a cells were treated with holotransferrin (HTF, 500 μg/mL) and imaged following the timing protocols 1 to 3 (Figure 2D, E, S3). HTF is an iron transport protein that delivers iron into cells after binding to its receptor on the cell membrane (Giometto et al., 1993). Treating cells with 500 μg/mL HTF enhanced the RF-induced activation of TRPV4^FeRIC^ using all protocols (Figure 2D, E; Table S1; No RF 24-h: 31 ± 9.5; RF 24-h: 520 ± 50.2; No RF 48-h: -0.5 ± 7.3; RF 48-h: 260.6 ± 52.3; No RF 72-h: -1.2 ± 4.8; RF 72-h: 550.1 ± 50). Notably, in TRPV4^FeRIC^-expressing N2a cells imaged 72-h after transfection, 500 μg/mL HTF treatment produced about a 10-fold increase in the RF-induced increase of GCaMP6 AUC relative to non-treated cells (Figure 2D, E; Table S1). Moreover, 500 μg/mL HTF also enabled RF at 465 kHz to activate TRPV4^FeRIC^ in cells imaged at 72-h post-transfection (Figure S2 D, Table S1; No RF + HTF 72- h: -1.2 ± 4.8 AUC; RF + HTF 72-h: 81.4 ± 19.4 AUC). Because the timing protocol 2 was less effective in activating TRPV4^FeRIC^ upon RF stimulation, we used only protocols 1 and 3 to evaluate the effects of other experimental variables.

Next, we examined the effects of different iron sources on the RF-induced activation of TRPV4^FeRIC^ using the timing protocol 1. Cells were supplemented with HTF, ferric citrate or a combination of HTF and ferric citrate. For control experiments, cells were supplemented with apotransferrin (apoTf) which is the iron-free version of HTF. In TRPV4^FeRIC^-expressing N2a cells, HTF enhanced the RF-induced Ca^2+^ responses at about 100 μg/mL; however, increasing HTF up to 2000 μg/mL did not enhance the RF responsiveness and produced vacuolation in several cells (Figure 2D-G, S3A). Treating the cells with apoTf at 100 μg/mL did not change the RF-induced Ca^2+^ responses with respect to untreated cells. Nevertheless, apoTf at higher concentrations enhanced RF-induced Ca^2+^ responses (Figure 2H, I). The unexpected effect of apoTf might be because, although it does not contain bound iron when added to cultured cells, a fraction of apoTf can bind the available iron from the culture medium becoming HTF and then delivering iron into cells. We also tested the effects of ferric citrate on RF-induced Ca^2+^ responses. At all examined concentrations (10, 50, and 100 μM), ferric citrate negatively impacted the RF-induced Ca^2+^ responses. Moreover, ferric citrate largely produced cell vacuolization (Figure S3B). Finally, in cells treated with a combination of suboptimal HTF (50 μg/mL) and ferric citrate (10 and 50 μM), RF stimulation activated robust Ca^2+^ responses in nearly all tested cells; although this treatment produced the highest observed responsiveness to RF stimulation, it also produced severe cell vacuolation (Figure S3C). Furthermore, treating the cells with 50 μg/mL HTF and 100 μg/mL ferric citrate μM abolished the RF-induced activation of TRPV4^FeRIC^ (Figure S3C).

These results indicate that increasing cellular iron import enhances the RF-induced Ca^2+^ responses in cells expressing TRPV4^FeRIC^. However, iron overload abolishes the magnetic activation of TRPV4^FeRIC^ and produces cell vacuolation. These results also indicate that the loss of RF responsiveness in TRPV4^FeRIC^-expressing N2a cells over time is associated with ferritin iron load. Nevertheless, previously we showed that expression of FeRIC channels for periods up to 72-h did not alter the cellular iron bioavailability (Hutson et al., 2017), suggesting that any potential progressive iron depletion, if it occurs, is limited to those ferritins coupled to FeRIC channels.

### IV. Abolishing the temperature sensitivity of TRPV4^FeRIC^ and lowering the extracellular Ca^2+^ levels prevent the functional downregulation of ferritin-tagged TRPV

Because some TRPV channels are constitutively active, cells prevent the cytotoxic effect of excessive Ca^2+^ influx by regulating their expression at the cell membrane (Bezzerides et al., 2004; Ferrandiz-Huertas et al., 2014; Montell, 2004; Planells-Cases and Ferrer-Montiel, 2007; Sanz-Salvador et al., 2012; Shukla et al., 2010). Diverse factors modulate the TRPV4 recycling from the cell membrane to cytosolic vesicles including temperature, continuous stimulation with agonists, extracellular Ca^2+^ levels, among others (Baratchi et al., 2019; Jin et al., 2011). Incubating cells at 37°C over extended periods of time may contribute to TRPV4 recycling. To examine if abolishing the temperature sensitivity of TRPV4^FeRIC^ prevents its functional downregulation, we used the temperature-insensitive mutant TRPV4^ΔTFeRIC^ (Y555A/S556A) (Duret et al., 2019; Hernández-Morales et al., 2020; Voets et al., 2002). RF stimulation significantly increased the GCaMP6 fluorescence in TRPV4^ΔTFeRIC^-expressing N2a cells imaged at either 24 or 72-h post-transfection compared to unstimulated cells (Figure 3A, B, table S1; No RF 24-h: 4.3 ± 2.2; RF 24-h: 393.8 ± 62.5; No RF 72-h: 7.8 ± 3.4; RF 72-h: 399.6 ± 42.4; p<0.0001) and this effect was inhibited with GSK219 (Figure 3A, B; Table S1). These results indicate that incubating the cells at 37°C for days likely affects the TRPV4^FeRIC^ functional expression. However, this effect should not be confused with that observed in Fig 1 where cells were incubated at 32°C and 37°C for about 20-30 min. The role of temperature in TRPV4^FeRIC^ functional expression at those different timescales, min versus days, might be different.

**Figure 3.**
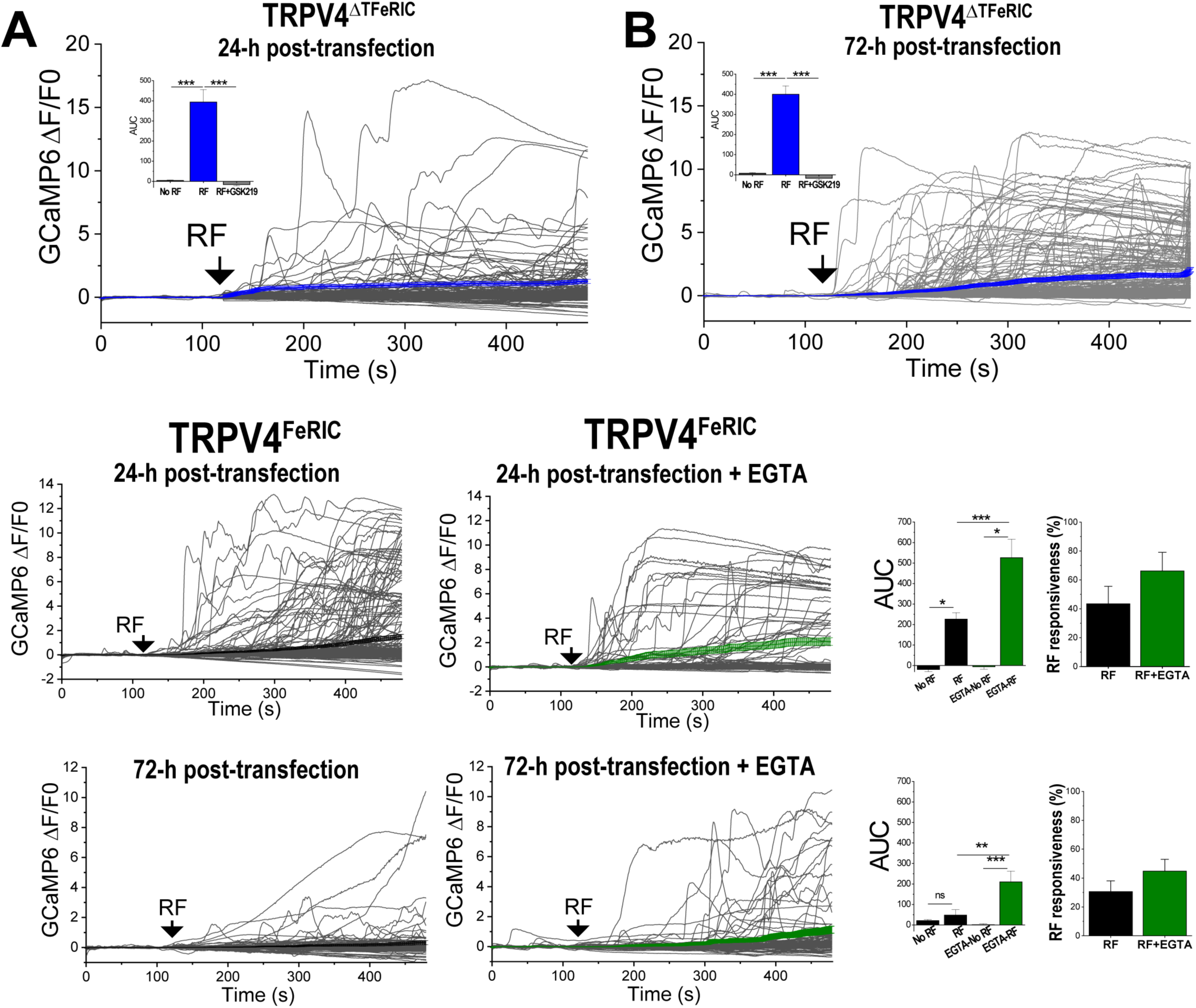
Temperature sensitivity and extracellular Ca^2+^ levels influence the RF activation efficiency of TRPV4^FeRIC^. (**A, B**) Average changes (± SEM) in GCaMP6 ΔF/F0 in N2a cells expressing TRPV4^ΔTFeRIC^ following continuous exposure to RF for 6 min (starting at black arrows). Cells were imaged (**A**) 24-h or (**B**) 72-h post-transfection. Insets: average changes (± SEM) in GCaMP6 AUC for the period of RF stimulation. (**C**) Average changes (± SEM) in GCaMP6 ΔF/F0 in N2a cells expressing TRPV4^FeRIC^ imaged 24-h post-transfection following RF stimulation. Right, cells were treated with EGTA after transfection. (**D**) Average changes (± SEM) in GCaMP6 AUC for the period of RF stimulation from experiments in (C). (**E**) Average changes (± SEM) in GCaMP6 ΔF/F0 in N2a cells expressing TRPV4^FeRIC^ imaged 72-h post-transfection following RF stimulation. Right, cells were treated with EGTA after transfection. (**F**) Average changes (± SEM) in GCaMP6 AUC for the period of RF stimulation from experiments in (E). Where applicable, either p<0.05 (*), p<0.001 (**), or p<0.0001 (***) was considered a statistically significant difference. See also Table S1.

Next, we examined if lowering the extracellular Ca^2+^ levels prevents the functional downregulation of TRPV4^FeRIC^, as it has been reported for TRPV1 (Sanz-Salvador et al., 2012). To lower the extracellular Ca^2+^ from ∼1.8 to ∼0.8 mM, the TRPV4^FeRIC^-expressing cells were incubated with culture medium supplemented with 1 mM EGTA. RF stimulation significantly increased the GCaMP6 fluorescence in EGTA-treated N2a cells expressing TRPV4^FeRIC^ imaged at either 24 or 72-h post-transfection with respect to unstimulated cells (Figure 3C-F; Table S1; No RF-EGTA 24-h: -5.5 ± 13; RF-EGTA 24-h: 526.2 ± 90.4; No RF-EGTA 72-h: 1.5 ± 5.4; RF-EGTA 72-h: 209.9 ± 52.8; p<0.0001). RF-induced increase in GCaMP6 fluorescence was larger in EGTA-treated TRPV4^FeRIC^-expressing N2a cells relative to untreated cells (Figure 3C-F, **insets**). Remarkably, EGTA treatment rescued the RF responsiveness in N2a cells expressing TRPV4^FeRIC^ for longer incubation periods (Figure 3E; Table S1).

These observations indicate that abolishing the temperature sensitivity of TRPV4^FeRIC^ or lowering the extracellular Ca^2+^ levels prevent its functional downregulation and consequent loss of RF responsiveness. Since the temperature-insensitive TRPV4^ΔTFeRIC^ is activated with RF but does not suffer downregulation, it can be an ideal candidate for use in *in vivo* applications of magnetogenetic techniques.

### **V.** Use of fluorescent Ca^2+^ dyes for monitoring RF-induced Ca^2+^ responses in cells expressing ferritin-tagged TRPV4

It has been reported that some Ca^2+^ dyes, such as Fura-2, may interfere with intracellular Ca^2+^ signaling (Alonso, 2003; Smith et al., 2018). Therefore, we asked if Fura-2 affects the RF-induced activation of TRPV4^FeRIC^. N2a cells expressing GCaMP6 plus TRPV4^FeRIC^ were loaded with Fura-2 AM (1 μM) (Figure S4A). Because the Fura-2 excitation/emission profile is in the UV wavelength range it does not interfere with GCaMP6 emission. Fura-2 abolished the RF-induced increase in GCaMP6 AUC in TRPV4^FeRIC^-expressing N2a cells imaged 24-h post transfection (Figure 4A, C; Table S1; RF: 550.1 ± 51; RF + Fura-2: -4.3 ± 3.9; p<0.0001). The inhibitory effect of Fura-2 was corroborated in TRPV4^FeRIC^-expressing N2a cells treated with 500 μg/mL HTF and imaged 72-h post-transfection, an experimental condition that gives a robust cytosolic Ca^2+^ increase upon RF stimulation (Figure 4B, C; Table S1; RF: 226.3 ± 31.4; RF + Fura-2: 2.8 ± 6; p<0.0001). To examine if other Ca^2+^ indicators inhibit the RF-induced activation of TRPV4^FeRIC^, next we monitored the cytosolic Ca^2+^ levels using the fluorescent dye Fluo-4 AM. In TRPV4^FeRIC^-expressing N2a cells imaged 24-h post-transfection and loaded with Fluo-4 (Figure S4B), RF stimulation significantly increased the Fluo-4 fluorescence relative to unstimulated cells (Figure 4D, F, S4; Table S1; Fluo-4 No RF: 9.4 ± 2.8; Fluo-4 RF: 215.3 ± 17.9; p<0.0001). Remarkably, using the protocol 1, the RF-induced Ca^2+^ responses had similar values when assessed with either GCaMP6 or Fluo-4 (Figure 4F). The RF-induced increase of Fluo-4 fluorescence was also corroborated in N2a cells expressing TRPV4^FeRIC^ treated with 500 μg/mL HTF and imaged 72-h post-transfection (Figure 4E, F; Table S1). These results indicate that, in TRPV4^FeRIC^-expressing N2a cells, Fura-2 inhibits the RF-induced cytosolic Ca^2+^ rise whereas Fluo-4 replicates those results observed using GCaMP6. These observations may guide the selection of the Ca^2+^ indicator for examining the RF-induced activation of ferritin-tagged TRPV.

**Figure 4.**
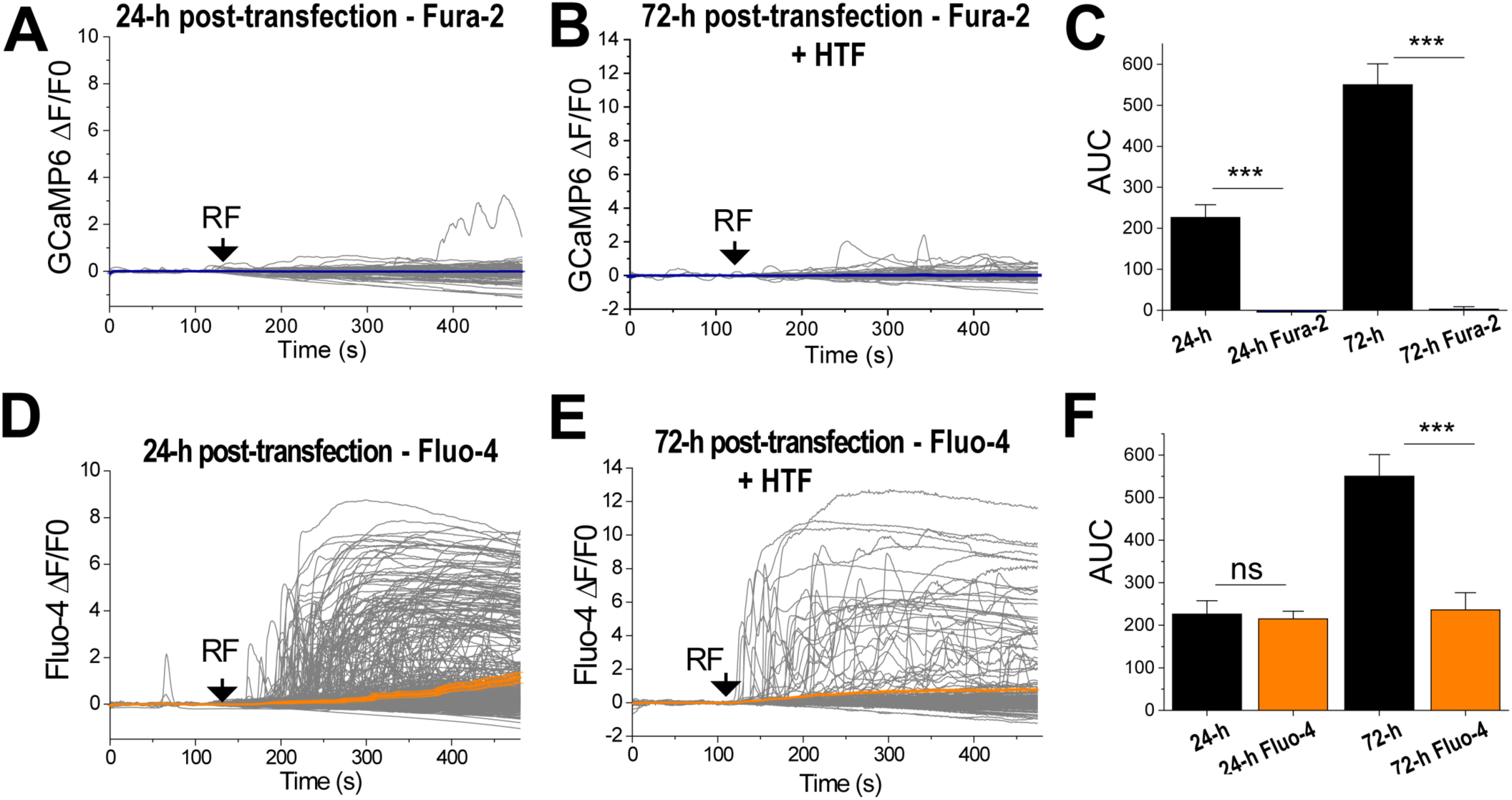
Fura-2 but not Fluo-4 decreased the ability of RF to activate TRPV4^FeRIC^. (**A, B**) Average changes (± SEM) in GCaMP6 ΔF/F0 in N2a cells expressing TRPV4^FeRIC^ following exposure to RF for 6 min (starting at black arrows). Cells were imaged (**A**) 24-h or (**B**) 72-h post-transfection and incubated 1-h before RF stimulation with Fura2 (1μM). Cells in (B) were supplemented with 500 μM HTF. (**C**) Average changes (± SEM) in GCaMP6 AUC in N2a cells expressing TRPV4^FeRIC^ in the absence (black bars) or the presence of Fura-2 (blue bars). (**D, E**) Average changes (± SEM) in Fluo-4 ΔF/F0 in N2a cells expressing TRPV4^FeRIC^ following exposure to RF for 6 min (starting at black arrows). Cells were imaged (**D**) 24-h or (**E**) 72-h post-transfection. Cells in (E) were supplemented with 500 μM HTF. Cells were loaded with 10 μM Fluo4 at 37°C for 1.5-h. (**F**) Average changes (± SEM) in GCaMP6 (black bars) and Fluo-4 (orange bars) AUC in N2a cells expressing TRPV4^FeRIC^ upon RF stimulation. Where applicable, either p<0.05 (*), p<0.001 (**), or p<0.0001 (***) was considered a statistically significant difference. See also Figure S4 and Table S1.

## Recommendations for *in vitro* protocols that activate transiently expressed TRPV^FeRIC^ with RF fields

Based on the observations described in this report, we have the following recommendations for FeRIC-based magnetogenetic techniques for RF-induced activation of cells in *in vitro* systems. For a detailed methodological description, please refer to the STAR Methods section.

1. Using our RF system, **RF fields at high MHz frequencies** are efficient in increasing the cytosolic Ca^2+^ levels in cells expressing ferritin-tagged TRPV.
2. In our experimental conditions, the **temperature-insensitive TRPV4^ΔTFeRIC^** produces a longer-lasting ability to remotely control cytosolic Ca^2+^ levels with RF fields than **TRPV4^FeRIC^**.
3. For monitoring RF-induced modulation of the cytosolic Ca^2+^ levels we recommend using genetically encoded Ca^2+^ indicators such as **GCaMP6**.
4. For seeding, transfecting, and examining the magnetic control of ferritin-tagged TRPV we recommend the **timing protocol 1**. In this protocol, cultured cells are allowed to transiently express the ferritin-tagged TRPV up to 24-h post-transfection.

i. It is important to optimize the expression levels of ferritin-tagged TRPV. **Overexpression** of TRPV4^FeRIC^ can cause cytotoxic effects.
ii. If longer expression periods are desired, then implement recommendations 5 and 6 of this list.
5. To increase iron loading of the ferritins coupled with TRPV, supplement the culture medium with **HTF or apoTf**. We observed that HTF at high concentration (above 1000 μg/mL) resulted in an appearance of intracellular vacuoles. It is recommended to determine the optimal concentration of HTF or apoTf for every specific *in vitro* system.
6. For those cases where it is not possible to allow for short time periods of ferritin-tagged TRPV expression or to supplement the culture medium with HTF or apoTf, the functional downregulation of those channels can be decreased by **lowering the extracellular Ca^2+^ levels**. This can be easily achieved by supplementing the culture medium with the Ca^2+^ chelator EGTA.

## Discussion

Because the main components of ferritin-based magnetogenetics, including ferritin and TRPV, are subjected to a diversity of cellular regulatory mechanisms, it is crucial to unify the experimental protocols to obtain reproducible results. The observations reported here may be used to guide the implementation of the current and future magnetogenetic tools.

Here we report that RF induces reproducible Ca^2+^ responses in cells expressing TRPV4^FeRIC^ at temperatures that are physiologically relevant. However, we acknowledge that the activation and decay kinetics of the RF-induced Ca^2+^ responses with FeRIC technology are in the hundreds of second scale. So, FeRIC technology is not yet suitable for addressing problems that require instantaneous or sub-second scale activation of cells.

In this study we found that the efficiency of FeRIC technology varies depending on several factors. We examined diverse experimental factors that affect the RF-induced activation of TRPV4^FeRIC^. These factors include the time period of TRPV4^FeRIC^ expression, cellular iron import, temperature sensitivity of TRPV4^FeRIC^, extracellular Ca^2+^ levels, and the use of different Ca^2+^ indicators. Promoting cellular iron import, mutating TRPV to abolish its temperature sensitivity or lowering extracellular Ca^2+^ levels enhance the RF-induced activation of TRPV4^FeRIC^. In contrast, longer time periods of TRPV4^FeRIC^ expression and Fura-2 inhibit the RF-induced activation of TRPV4^FeRIC^. Moreover, RF fields at high MHz frequencies is a stimulus that consistently activates TRPV4^FeRIC^ without the inconvenience of heating up the imaging saline solution.

Interestingly, longer periods of TRPV4^FeRIC^ expression drastically decrease the ability of RF to produce a Ca^2+^ rise in cells expressing those channels. The possible explanations are that cellular ferritins are progressively iron-depleted, the TRPV4^FeRIC^ are functionally downregulated, the coupling between ferritins and TRPV4^FeRIC^ is lost, or the sum of all these factors. We have previously reported that, at the maximum period of TRPV4^FeRIC^ expression tested (72-h post-transfection), the main iron-regulatory proteins mRNA are not altered by the expression of FeRIC channels (Hutson et al., 2017). This indicates that the cellular iron homeostasis is not disturbed in the FeRIC system, but maybe those ferritins coupled to FeRIC channels are iron depleted. The role of the ferritin iron load in the loss of RF responsiveness was corroborated in cells treated with HTF, apoTf or a combination of ferric citrate plus HTF. Increasing the cellular iron import may increase the ferritin iron load (Brier et al., 2020) and consequently enhances the RF-induced activation of TRPV4^FeRIC^. However, it is necessary to optimize the iron source (HTF, apoTf, ferric citrate) and the concentration needed for each specific experimental model to prevent cytotoxicity. While in HEK293T cells expressing anti-GFP–TRPV1/GFP–ferritin the optimal iron supplementation was a combination of 2 mg/mL HTF and 500 μM ferric citrate (Brier et al., 2020), N2a cells expressing TRPV4^FeRIC^ needed a 20 times lower concentration of HTF. Moreover, ferric citrate as low as 10 μM was cytotoxic (Figure S3). These differences could be because cells expressing anti-GFP–TRPV1/GFP–ferritin overexpress ferritin which might help them to handle the iron excess. In contrast, in FeRIC technology, cells are not overexpressing ferritin, but instead TRPV channels are coupled with endogenous ferritins. Then, excessive iron might overpass the cells’ capabilities to maintain iron homeostasis. Our results are consistent with reports about cell death caused by iron treatment at concentrations similar to what we used in our experiments (Baba et al., 2018; Chen et al., 2020; Lee et al., 2021).

TRPV channels are crucial components of most of the magnetogenetics approaches (Chen et al., 2015; Duret et al., 2019; Hernández-Morales et al., 2020; Huang et al., 2010; Hutson et al., 2017; Stanley et al., 2015, 2016; Wheeler et al., 2016). Notably, the functional expression of TRPV is regulated by activity-dependent mechanisms, resulting in dynamic trafficking between the cell membrane and the cytosolic vesicle pool. It has been shown that TRPV channels are maintained in vesicles and their activators cause their incorporation into the cell membrane via exocytosis (Baratchi et al., 2019; Bezzerides et al., 2004; Montell, 2004; Planells-Cases and Ferrer-Montiel, 2007). However, either constitutive or continuous activation of TRPV1 and TRPV4 causes their translocation from the cell membrane to recycling vesicles to prevent the cytotoxicity effects of excessive Ca^2+^ influx (Baratchi et al., 2019; Jin et al., 2011; Sanz-Salvador et al., 2012; Shukla et al., 2010; Tian et al., 2019). In our FeRIC system, cells expressing TRPV4^FeRIC^ are incubated at 37°C, a temperature above the TRPV4 activation threshold (34°C) (Huang et al., 2018; Liedtke et al., 2000; Strotmann et al., 2000). At this temperature, TRPV4^FeRIC^ is likely constitutively and continuously activated resulting in its functional downregulation, an effect more pronounced with longer periods of TRPV4^FeRIC^ expression. The role of the temperature-mediated downregulation was corroborated with the temperature-insensitive TRPV4^ΔTFeRIC^ whose RF responsiveness was not disturbed by longer periods of expression (Figure 3A, B). However, despite the fact that temperature may play a role in the functional expression of TRPV4^FeRIC^, here we demonstrated that these channels are still activated with RF at physiologically relevant temperatures (Figure 1F, H). The functional expression of TRPV4 is also modulated by extracellular Ca^2+^ levels (Baratchi et al., 2019; Jin et al., 2011). Consistent with this finding, in our FeRIC system, lowering the extracellular Ca^2+^ levels enhanced the RF responsiveness of TRPV4^FeRIC^ (Figure 3C-F). Therefore, our observations indicate that abolishing the temperature sensitivity of TRPV or lowering the extracellular Ca^2+^ levels prevent the functional downregulation of ferritin-tagged TRPV.

Another unexpected finding was the inhibitory effect of Fura-2 on the RF-induced cytosolic Ca^2+^ rise in N2a cells expressing TRPV4^FeRIC^ (Figure 4A-C). It has been reported that Fura-2 interferes with some intracellular Ca^2+^ signaling pathways (Alonso, 2003). Additionally, it has also been reported that Fluo-4 affects cell functions; nonetheless, we observed similar results using either GCaMP6 or Fluo-4 as the Ca^2+^ indicator (Figure 4D-F). Therefore, it might be relevant to select a suitable Ca^2+^ indicator when examining the cytosolic Ca^2+^ rise produced by the magnetic activation of ferritin-tagged TRPV.

While several independent studies have reported the magnetic control of TRPV1 and TRPV4 (Brier et al., 2020; Duret et al., 2019; Hernández-Morales et al., 2020; Hutson et al., 2017; Stanley et al., 2015, 2016; Wheeler et al., 2016), magnetogenetics has generated controversies about its underlying mechanisms and experimental reproducibility. Our study indicates that several experimental factors can negatively affect the efficacy of RF fields in activating ferritin-tagged TRPV4. The observations reported here set a precedent for examining diverse experimental factors that might be involved in some inconsistences among different magnetogenetic studies.

In conclusion, our study identified several experimental factors that impact the efficacy of RF fields on inducing cytosolic Ca^2+^ rise in cells expressing ferritin-tagged TRPV. Our findings pointed out that ferritin-based magnetogenetics are sensitive to diverse experimental factors that may disturb the functional expression and function of ferritin and/or TRPV. This study offers a guide for optimizing the current ferritin-based magnetogenetic tools and protocols to obtain reproducible results.

## Limitations of the study

This study is limited to a single ferritin-based magnetogenetic technique called FeRIC. Further studies are needed to corroborate our findings in other magnetogenetic approaches that use, for example, cells stably expressing the ferritin-tagged TRPV or chimeric ferritins. Furthermore, it is necessary to examine the diverse experimental factors that could impact the efficacy of RF fields in activating ferritin-tagged TRPV4 in in vivo models.

## STAR methods

### Resource availability

#### Lead contact

Further information and requests for resources and reagents should be directed to and will fulfilled by the Lead Contact, Chunlei Liu.

#### Materials availability

Plasmids will be made available upon request and will require a material transfer agreement from C.L.

#### Data and code availability

- The Ca^2+^ data analysis code used during this study is available at GitHub: https://github.com/LiuCLab/FeRIC/blob/master/FeRIC_Ca_imaging.m.
- The Magnetic/electric field simulation code used during this study is available at GitHub: https://github.com/LiuCLab/FeRIC/blob/master/FeRIC_FDTD_simulation.m.
- The cDNA sequence of TRPV4^FeRIC^ is available at GenBank with the identifier TRPV4FeRIC: MT025942 (https://www.ncbi.nlm.nih.gov/nuccore/).
- The Ca^2+^ imaging analysis files are available at Mendeley Data (doi: 10.17632/k9tfrgmjzg.1).
- The dataset identifiers and accession numbers are in the key resources table.
- Any additional information required to reanalyze the data reported in this paper is available from the lead contact upon request.

### Experimental Model and Subject Details

#### Cell lines

Neuro2a cell line (N2a, ATC, CCL-13) was used. N2a cells were obtained from the UCB Cell Culture Facility (University of California Berkeley). Cell identity and negative Mycoplasma contamination were verified by the UCB Cell Culture Facility. Cells were maintained in Dulbeccos’s Modified Eagle Medium (DMEM, Gibco) supplemented with 10% fetal bovine serum (FBS, hyclone) and 100 units/mL penicillin, and 100 mg/mL streptomycin at 37°C and 5% CO2. For described experiments, cell lines were employed between the passages 5 to 20.

### Method details

#### Plasmids

The constructs TRPV4^FeRIC^, TRPV4^ΔTFeRIC^, and TRPV4^WT^ were obtained as described previously (Hutson et al., 2017; Hernandez-Morales et al., 2020). To generate the TRPV4^WT^ construct, we used full-length rat TRPV4 cDNA, which was a gift from R. Lefkowitz (Duke University). Spe I and Not I restriction sites were introduced using PCR. The full-length wild-type TRPV4 was subcloned into the PLVX-IRES-mCherry vector to generate TRPV4^WT^ (Clontech, Catalog No. 631237). To generate the TRPV4^FeRIC^ construct, PCR primers were designed to eliminate the 3′ stop site in wild-type TRPV4 and introduce a 3′ Not I site. PCR primers introducing a 5′ Not I site, and a 3′ Bam HI site and a stop codon were used to amplify human Kininogen1 domain 5 (FeRIC). This FeRIC fragment was subcloned into the Xba I and BamH 1 sites within the PLVX-IRES-mCherry vector containing TRPV4. All completed constructs were sequence-verified by the Molecular Cell Biology Sequencing Facility (UC Berkeley) and analyzed using MacVector 13.0 and Serial Cloner 2.6.

#### RF coils

For RF at the 180 MHz range, one RF emitting coil with diameter of 5 cm was used. The RF coil was made of two loops of wire. For RF at 465 kHz, one coil with a diameter of 5 cm and made of 7 loops of wire was used. The coils were connected in series with tuning capacitors forming an LC circuit, and were tuned to a resonance frequency of about 180 MHz or 465 kHz. They were then matched to 50 ohms using parallel capacitors. For 180 MHz, the RF signal was generated by a broadband (35 MHz to 4.4 GHz) signal generator (RF Pro Touch, Red Oak Canyon LLC) and amplified using a 5W amplifier (Amplifier Research, model 5U100, 500 kHz to 1000 MHz). For 465 kHz, the RF signal was generated by a DDS signal generator and amplified using a custom-built 50W RF amplifier (operational from 100 kHz to 3 MHz). The magnetic field produced by the coils was measured using EMC near-field probes (Beehive Electronics model 100B and TekBox TBPS01 H5 probe) connected to a spectrum analyzer (RSA3015E-TG, Rigol Technologies). The Beehive 100B probe was provided with magnetic field calibration data from the manufacturer and proper scaling factors for 180 MHz and 465 kHz measurements were interpolated based on this data. The Rigol spectrum analyzer was manufacturer calibrated with a system registered to ISO 9001:2008. The TekBox probe was not calibrated. Using the Beehive probe and the Rigol spectrum analyzer, the magnetic field strength was measured to be about 1.6 µT for 180 MHz or 31 µT for 465 kHz at a location about 3 mm above the cell culture dishes. These are the values used throughout the manuscript. The measurements were made slightly above the cell culture dish location because the probes could not be lowered into the coils without changing the angle of the probes with respect to the magnetic field. Simulations, however, predict that there is only about a 3% decrease in magnetic field strength 3 mm above the center of the coil. Using the TekBox probe and the Rigol spectrum analyzer, the magnetic field strength was measured to be about 2.3 µT for 180 MHz or 46 µT for 465 kHz. These values are provided as a validation and for potential TekBox probe users’ convenience.

#### Chemical transfection – Lipofectamine LTX Plus reagent

N2a cells were plated on non-coated glass-bottom 35-mm dishes. Cells were cultured in DMEM supplemented medium. After 18-24 hours, cells were transfected using the Lipofectamine LTX Plus reagent (ThermoFisher #15338030) with GCaMP6 (GCaMP6 medium, Addgene cat.40754) and TRPV4^FeRIC^ channels. For transfection, OptiMEM free serum medium (ThermoFisher #31985088) was used to prepare the DNA/Lipofectamine LTX mix. Transfection mix had the following composition per each 35-mm dish: 300 μL OptiMEM, 4 μL Lipofectamine LTX, 3 μL PLUS reagent, 0.7 μg TRPV4 DNA, and 0.7 μg of GCaMP6 DNA. Where indicated, EGTA (pH 8, ThermoFisher Scientific, #NC0997810) was added to dishes after transfection to achieve 1 mM final concentration. It was estimated that 1 mM of EGTA decreases the Ca^2+^ concentration in the DMEM culture medium from ∼1.8 to ∼0.8 mM (https://somapp.ucdmc.ucdavis.edu/pharmacology/bers/maxchelator/CaEGTA-TS.htm).

#### Protocols for seeding, transfection, and HTF treatment

##### Timing protocol 1 (24-h post-transfection)

N2a cells were plated on non-coated glass-bottom 35-mm dishes. Cells were cultured in DMEM supplemented medium. After 20-24 hours, cells were transfected using the Lipofectamine LTX Plus reagent (ThermoFisher #15338030) with GCaMP6 (GCaMP6 medium, Addgene cat.40754) and FeRIC channels as described above. Ca^2+^ imaging experiments were conducted at 20-24 h post-transfection.

##### Timing protocol 1 (24-h post-transfection) – HTF and apoTf treatment

Where indicated, HTF or apoTf were added to culture medium immediately after transfection to achieve 25 to 2000 μg/mL final concentration. Ca^2+^ imaging experiments were conducted at 20-24 h post-transfection.

##### Timing protocol 1 (24-h post-transfection) – ferric citrate treatment

Where indicated, ferric citrate (10 to 100 μM) was added to culture medium at 6-h post-transfection. Adding ferric citrate immediately after transfection prevents the expression of both TRPV4^FeRIC^ and GCaMP6. At 20-24 h post-transfection, Ca^2+^ imaging experiments were conducted.

##### Timing protocol 2 (48-h post-transfection)

N2a cells were seeded on 25-cm flask and cultured in DMEM supplemented medium. After 20-24 hours (day 1), cells were transfected using the Lipofectamine LTX Plus reagent (ThermoFisher #15338030) with GCaMP6 (GCaMP6 medium, Addgene cat.40754) and TRPV4^FeRIC^. For transfection, OptiMEM free serum medium (ThermoFisher #31985088) was used to prepare the DNA/Lipofectamine LTX mix. Transfection mix had the following composition per 25-cm flask: 900 μL OptiMEM, 12 μL Lipofectamine LTX, 9 μL PLUS reagent, 2.1 μg TRPV DNA, and 2.1 μg of GCaMP6 DNA. At 24-h post-transfection, N2a cells were plated on non-coated glass-bottom 35-mm. Next, Ca^2+^ imaging experiments were conducted at 48-h post-transfection.

##### Timing protocol 2 (48-h post-transfection) – HTF treatment

Where indicated, HTF was added to culture medium immediately after transfection to achieve 500 μg/mL final concentration. HTF treatment was maintained until Ca^2+^ imaging experiments were conducted.

##### Timing protocol 3 (72-h post-transfection)

N2a cells were plated on non-coated glass-bottom 35-mm dishes at very low density (1 x10E5 cells). Cells were cultured in DMEM supplemented medium. After 20-24 hours, cells were transfected using the Lipofectamine LTX Plus reagent (ThermoFisher #15338030) with GCaMP6 (GCaMP6 medium, Addgene cat.40754) and TRPV4^FeRIC^ as described above. At 24-h post-transfection, the culture medium was replaced with fresh culture medium. Ca^2+^ imaging experiments were conducted at 72-h post-transfection.

##### Timing protocol 3 (72-h post-transfection) – HTF treatment

N2a cells were plated on non-coated glass-bottom 35-mm dishes at very low density (1 x10E5 cells). Cells were cultured in DMEM supplemented medium. After 20-24 hours, cells were transfected using the Lipofectamine LTX Plus reagent (ThermoFisher #15338030) with GCaMP6 (GCaMP6 medium, Addgene cat.40754) and TRPV4^FeRIC^ as described above. At 24-h post-transfection, the culture medium was replaced with fresh culture medium supplemented with HTF (500 μg/mL). Ca^2+^ imaging experiments were conducted at 72-h post-transfection.

##### Ca^2+^ imaging – GCaMP6

Epifluorescence imaging experiments were conducted as previously described (Hutson et al., 2017). Cytosolic levels of Ca^2+^ were monitored by fluorescence imaging of cells positive for GCaMP6. Cells expressing TRPV4^FeRIC^ channels were identified as those cells with mCherry^+^ expression. Experiments were conducted using an upright AxioExaminer Z-1 (Zeiss) equipped with a camera (Axiocam 702 mono) controlled by Zen 2.6 software. Excitation light was delivered from a LED unit (33 W/cm2; Colibri 7 Type RGB-UV, Zeiss). mCherry was excited at 590/27 nm and emission was collected at 620/60 nm. GCaMP6 was excited at 469/38 nm and emission was collected at 525/50 nm. Illumination parameters were adjusted to prevent overexposure and minimize GCaMP6 photobleaching. All the experiments corresponding to a series were done under the same illumination settings. Images were captured with a W “Plan-Apochromat” 20x/1.0 DIC D=0.17 M27 75mm lens at 1 frame/s. Experiments were carried out at room temperature (20 -22 °C) using HBSS (Invitrogen, 14025092) that contains (in mM) 140 NaCl, 2.5 KCl, 1 MgCl2, 1.8 CaCl2, 20 HEPES, pH 7.3. HBSS was supplemented with 10 mM D-glucose. At the beginning of imaging experiments, a 35-mm glass-bottom dish containing the cells was washed three times with 1 mL of the HBSS. Next, the dish was placed onto the microscope stage and the cells were localized with transmitted illumination (bright-field). Next, with reflected illumination, the fluorescence signals from mCherry and GCaMP6 were corroborated and the field of interest was selected. Preferred fields were those with isolated and healthy cells. A thermocouple coupled to a thermistor readout (TC-344C, Warner Instruments) was placed inside the plate in direct contact with the HBSS. The temperature of the HBSS was monitored during the experiment (Temperature initial: 22.09°C; Temperature final: 22.03°C; ΔT: -0.06°C; n=305). Cells were rested for about 10 minutes before imaging. RF was delivered using a custom-built RF-emitting coil that fits the 35-mm tissue culture dish, as described above. Cells were stimulated with RF fields at 180 MHz (at 1.6 µT) or 465 kHz (at 31 µT). For each experiment, cells were imaged for the first 60-120 s with no stimulus (Basal) and followed with RF exposure for 4-6 min. After RF stimulation, cells were exposed to GSK101, which is a TRPV agonist, by pipetting 1 mL of the GSK101 diluted in HBSS (at 2μM) to reach the final 1μM concentration.

##### Ca^2+^ imaging – GCaMP6 plus Fura-2

For Fura-2 experiments, cells expressing GCaMP6 and TRPV4^FeRIC^ were loaded with Fura-2. Cells were incubated with Fura-2 AM (ThermoFisher scientific, F1221) at 1 μM for 30 min at room temperature in darkness. Next, cells were washed three times with HBSS and rested for 30 min at room temperature in darkness. To corroborate that cells were loaded with Fura-2, cells were imaged with a 365 nm excitation wavelength and emission was collected at 445/450 nm (Figure S4). Next, cytosolic levels of Ca^2+^ were monitored by fluorescence imaging of cells positive for GCaMP6 as described in the Ca^2+^ imaging – GCaMP6 section.

##### Ca^2+^ imaging – Fluo-4

Cytosolic levels of Ca^2+^ were monitored by fluorescence imaging of cells loaded with Fluo-4. Cells expressing TRPV4^FeRIC^ were incubated with Fluo-4 AM (ThermoFisher scientific, F14201) at 10 μM for 1.5 h at room temperature in darkness. Next, cells were washed three times with HBSS and rested for 30 min at room temperature in darkness. Fluo-4 was excited at 469/38 nm and emission was collected at 525/50 nm. Illumination parameters were adjusted to prevent overexposure and minimize Fluo-4 photobleaching. All the experiments corresponding to a series were done under the same illumination settings. Images were captured with a W “Plan-Apochromat” 20x/1.0 DIC D=0.17 M27 75mm lens at 1 frame/s. Next, cytosolic levels of Ca^2+^ were monitored by fluorescence imaging of cells loaded with Fluo-4 as described in the Ca^2+^ imaging – GCaMP6 section.

##### Yo Pro 1 assay

N2a cells were plated on non-coated glass-bottom 35-mm dishes. Cells were cultured in DMEM supplemented medium. After 18-24 hours, cells were transfected using the Lipofectamine LTX Plus reagent (ThermoFisher # 15338030) TRPV4^FeRIC^ or TRPV4^WT^. For transfection, OptiMEM free serum medium (ThermoFisher # 31985088) was used to prepare the DNA/Lipofectamine LTX mix. Transfection mix had the following composition per each 35-mm dish: 300 μL OptiMEM, 4 μL Lipofectamine LTX, 3 μL PLUS reagent, 0.7 μg TRPV4^FeRIC^ or TRPV4^WT^ DNA. After 24-hours of transfection, cells were washed three times with HBSS. Cell nuclei were stained with Hoechst 33342 (ThermoFisher Scientific, NucBlue® Live ReadyProbes® Reagent, R37605) by incubating the cells for 10 min at 37°C. Next, cells were treated with GSK101 (50 nM) plus Yo Pro 1 (1 μM) for 20 min at 37°C. Cells were washed three times with HBSS and imaged. mCherry was excited at 590/27 nm and emission was collected at 620/60 nm. Hoechst 33342 was excited at 365 nm and emission was collected at 445/50 nm. Yo Pro 1 was excited at 469/38 nm and emission was collected at 525/50 nm. For each independent experiment, cells were localized with transmitted illumination (bright-field) and 2-5 fields of view were randomly selected. Cells were imaged to observe mCherry, Yo Pro 1, and Hoechst 33342 fluorescence. All the experiments were done under the same illumination settings.

#### Quantification and statistical analysis

##### Simulations of the magnetic and electric fields produced by RF

The distribution of the electric (E) and magnetic (B) fields applied to the cells were simulated using the finite-difference time-domain (FDTD) method implemented by the openEMS project (Liebig et al., 2013) (https://openems.de/start/). The simulations were done considering the 5 cm-diameter RF coil containing the 3.5 cm-diameter dish half-filled with imaging saline solution (dish height: 1 cm, saline solution height: 0.5 cm). The dielectric constant (80) and conductivity (1.5 S/m) for the imaging saline solution was obtained from the literature (Davis et al., 2020). In the simulation, a signal with a center frequency of 180 MHz or 465 kHz was injected across the wires of a two-turn solenoid. The resulting time-domain fields were Fourier transformed, and the magnitudes of the B (T) and E (V/m) field vectors at each spatial location were calculated for 180 MHz or 465 kHz.

##### Ca^2+^ imaging analysis

Cytosolic Ca^2+^ levels were monitored in cells expressing GCaMP6 and either FeRIC or wild-type TRPV channels. The fluorescence intensity of GCaMP6 was acquired at 1 Hz. GCaMP6 fluorescence was computed in a cell-based analysis with a customized MATLAB (Release 2018b, MathWorks Inc., Natick, Massachusetts) code. The maximum intensity projection was performed along the time axis to get the maximum intensity signal of cells expressing GCaMP6 in the field of view. In experiments where cells were stimulated with agonists, the maximum intensity projection corresponds to those cells which expressed functional ion channels. The watershed algorithm (MATLAB implemented function: Watershed transform) (Meyer, 1994) was used to identify and label the cells to generate a cell-based mask for each experiment. The algorithm does not contain the motion correction component because the spatial movement of cells during the time-lapse acquisition was negligible. The GCaMP6 fluorescence intensity was measured for each masked-cell of the time-lapse acquisition (600 s). GCaMP6 fluorescence signal is presented as ΔF/F0, where F0 is the basal fluorescence averaged over 1-121 s before the start of stimulation and ΔF is the change in fluorescence over basal values. For analysis, the GCaMP6 fluorescence measurements corresponding to the first 5 frames were discarded because of the appearance of an inconsistent artifact. For each masked-cell, the data from 6-121 s were fit with a mono-exponential curve [f(t; a, b) = a*exp(b*t)]. The fitted curve was used to correct the GCaMP6 photobleaching effect over the entire acquisition period. The masked-cells that showed abnormal behavior, observed as the value for the growth factor (b) above 0.002, were excluded from the analysis. Masked-cells were considered responsive when the averaged ΔF/F0 over time during the period from 122 to 480 s (RF-stimulation) increased 10 times over the standard deviation of the GCaMP6 ΔF/F0 of the basal session. For each experimental group, the change in GCaMP6 ΔF/F0 and the corresponding AUC (t = 121 – 480 s) of all analyzed masked-cells, that corresponded to all TRPV-expressing cells, were averaged. The plots of the GCaMP6 ΔF/F0 changes correspond to the data obtained from all identified cells, including both responsive and non-responsive cells. The responsiveness was estimated for each independent experiment as the ratio of the responsive masked-cells relative to the total of analyzed masked-cells. For statistical analysis, the AUC (t = 122 – 480 s) was estimated for each masked-cell. Data are presented as the AUC of GCaMP6 fluorescence (± SEM), the fold change (± SEM) of the AUC of GCaMP6 fluorescence and responsiveness. The fold change was calculated by dividing the value of the AUC of GCaMP6 ΔF/F0 in the presence of the stimulus (RF) by the value of that in the absence of stimulation (No stimulus or No RF). In the case fold change is less than one, the reciprocal is listed (minus-fold change). For each experimental condition, more than 3 independent experiments were conducted. The MATLAB code that we used for the Ca^2+^ imaging analysis is available in Supplementary Materials.

##### Yo Pro 1 assay analysis

The number of total cells (Hoechst 33342 stained cell nucleus) and the number of Yo Pro 1 positive cells in a field of view was computed in a cell-based analysis with a customized MATLAB (Release 2018b, MathWorks Inc., Natick, Massachusetts) code.

##### Quantification and Statistical Analysis

All experiments were repeated a minimum of three times. Differences in continuous data sets were analyzed using Microcal OriginPro 2020 software (OriginLab). The hypothesis testing was performed using a one-way ANOVA, followed by Tukey’s post hoc test. When only two experimental groups were compared, the statistical probe applied was Student’s t-test. Data are means ± SEM. Where applicable, *p<0.05, **p<0.001, or ***p<0.0001 was considered a statistically significant difference.

## Acknowledgements

We thank Jingjia Chen (University of California, Berkeley) for coding the customized MATLAB code for counting the number of cells for the Yo Pro 1 experiments. We thank Koyam Morales-Weil for conducting control experiments that demonstrate that RF does not affect endogenous channels in N2a cells.

Research reported in this publication was in part supported by the National Institute of Neurological Disorders and Stroke of the National Institutes of Health under Award Number R01NS110554. The content is solely the responsibility of the authors and does not necessarily represent the official views of the National Institutes of Health.

## Author contributions

C.L. and M.H.-M. designed the project. M.H.-M. designed and conducted the experiments and analyses. V.H. built the RF coils and estimated the magnetic and electric field distribution in the experimental system. R.H.K. contributed to the Yo Pro experiments. M.H.-M. wrote the manuscript. C.L., V.H. and R.H.K. revised and edited the manuscript. All authors approved the manuscript.

## Inclusion and Diversity

One or more of the authors of this paper self-identifies as an underrepresented ethnic minority in science.

## Supplemental Figures and Table

**Figure S1.**
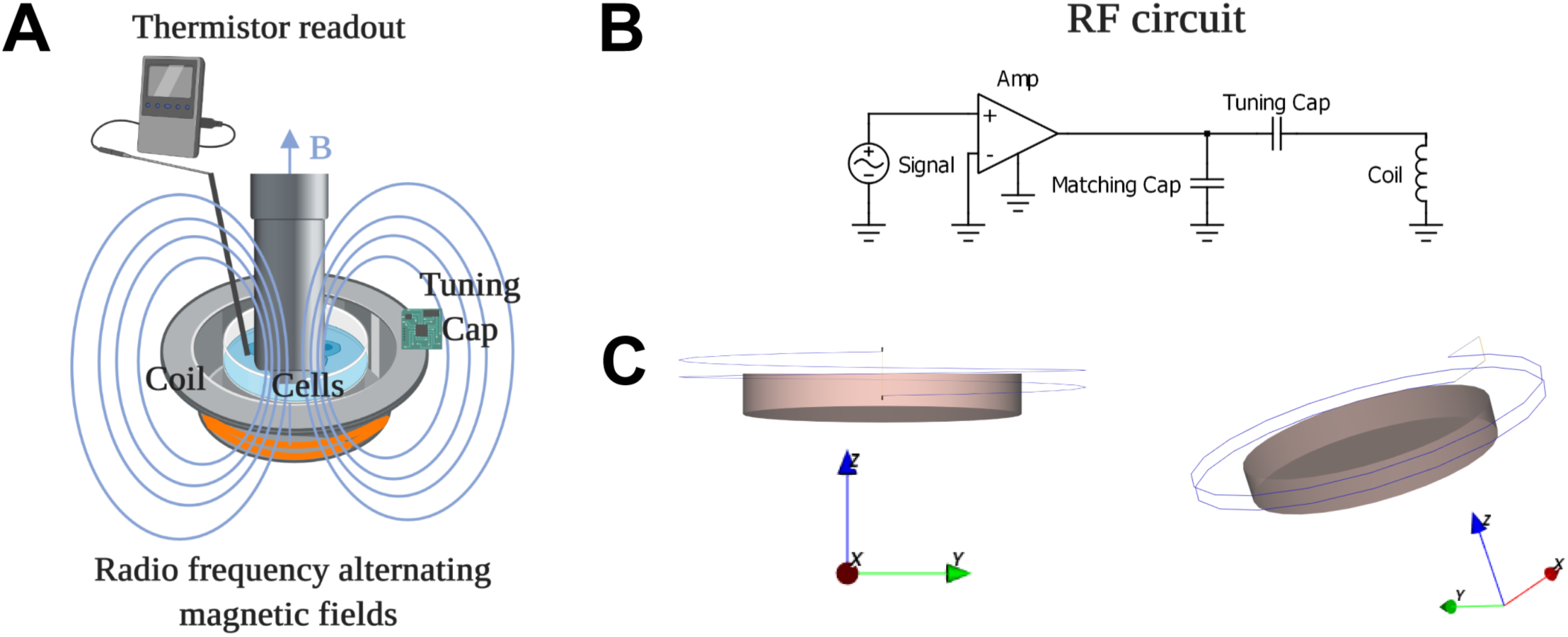
Scheme of the RF setup. Related to Figure 1. (**A**) Illustration of the experimental set up for RF stimulation of cultured cells. (**B**) Simplified RF circuit. (**C**) Two orientations of 3D models of the culture dish and the solenoid RF coil.

**Figure S2.**
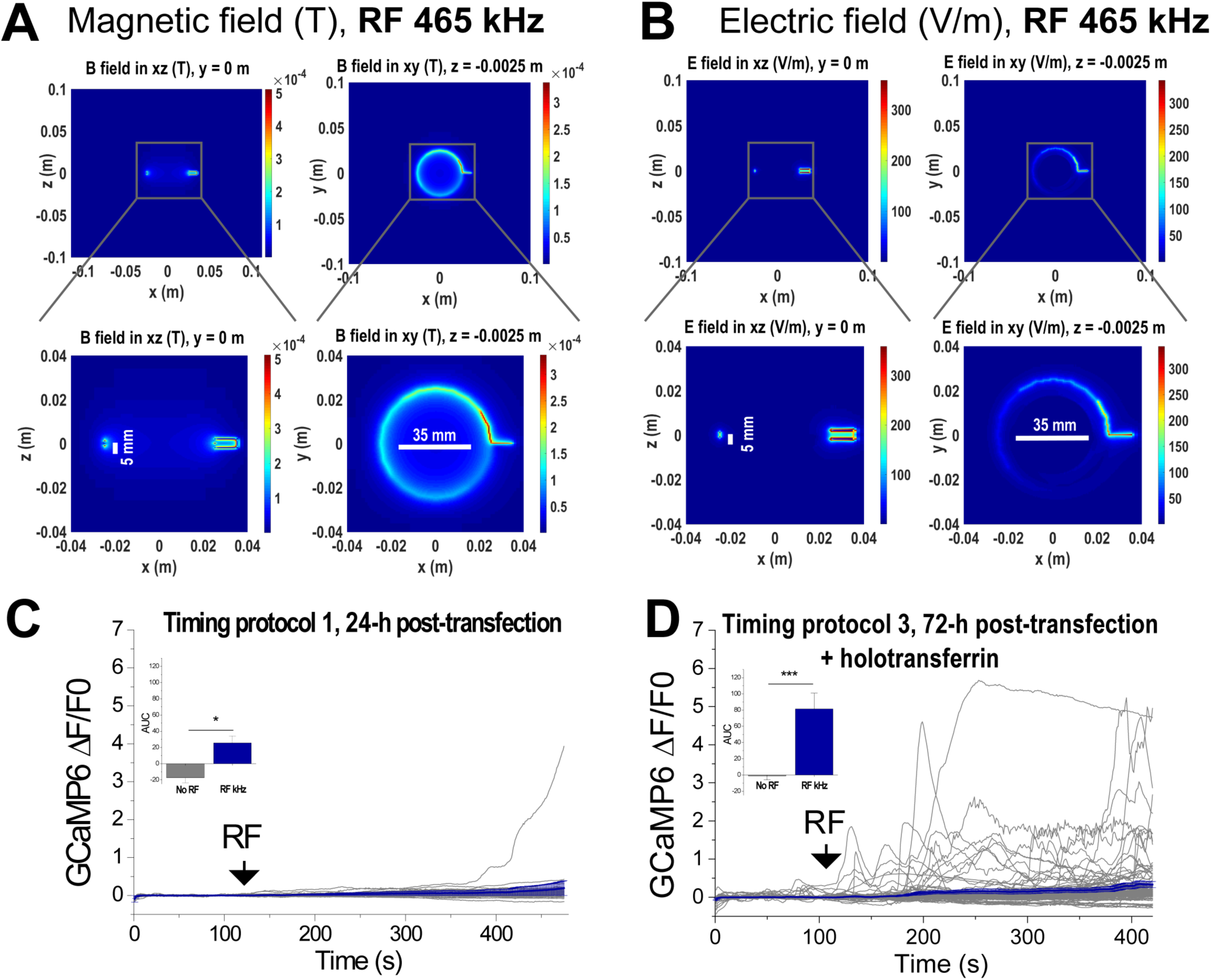
RF stimulation at kHz frequency at the μT range has limited efficiency for activating TRPV4^FeRIC^. Related to Figures 1 and 2. (**A**, **B**) Simulations of the magnetic and electric fields produced by RF at kHz. Distribution of the (**A**) magnetic and (**B**) electric fields in the culture dish produced by RF fields at 465 kHz and 31 µT; bottom: zoom-in of the culture dish. Vertical lines correspond to the 5mm-height of the saline solution being in the lower half of the coil. Horizontal lines: 35 mm-diameter of the culture dish. Cultured cells are at the center of the culture dish. (**C, D**) All changes and average changes (± SEM) in GCaMP6 ΔF/F0 in N2a cells expressing GCaMP6 plus TRPV4^FeRIC^ following stimulation with RF at 465 kHz and 31 μT for 6 min (black arrows). Cells were imaged (**C**) 24-h or (**D**) 72-h post-transfection. Cells in (D) were supplemented with 500 µM HTF after transfection. Insets: average changes (± SEM) in GCaMP6 AUC for the period of RF stimulation. Where applicable, either p<0.05 (*), p<0.001 (**), or p<0.0001 (***) was considered a statistically significant difference.

**Figure S3.**
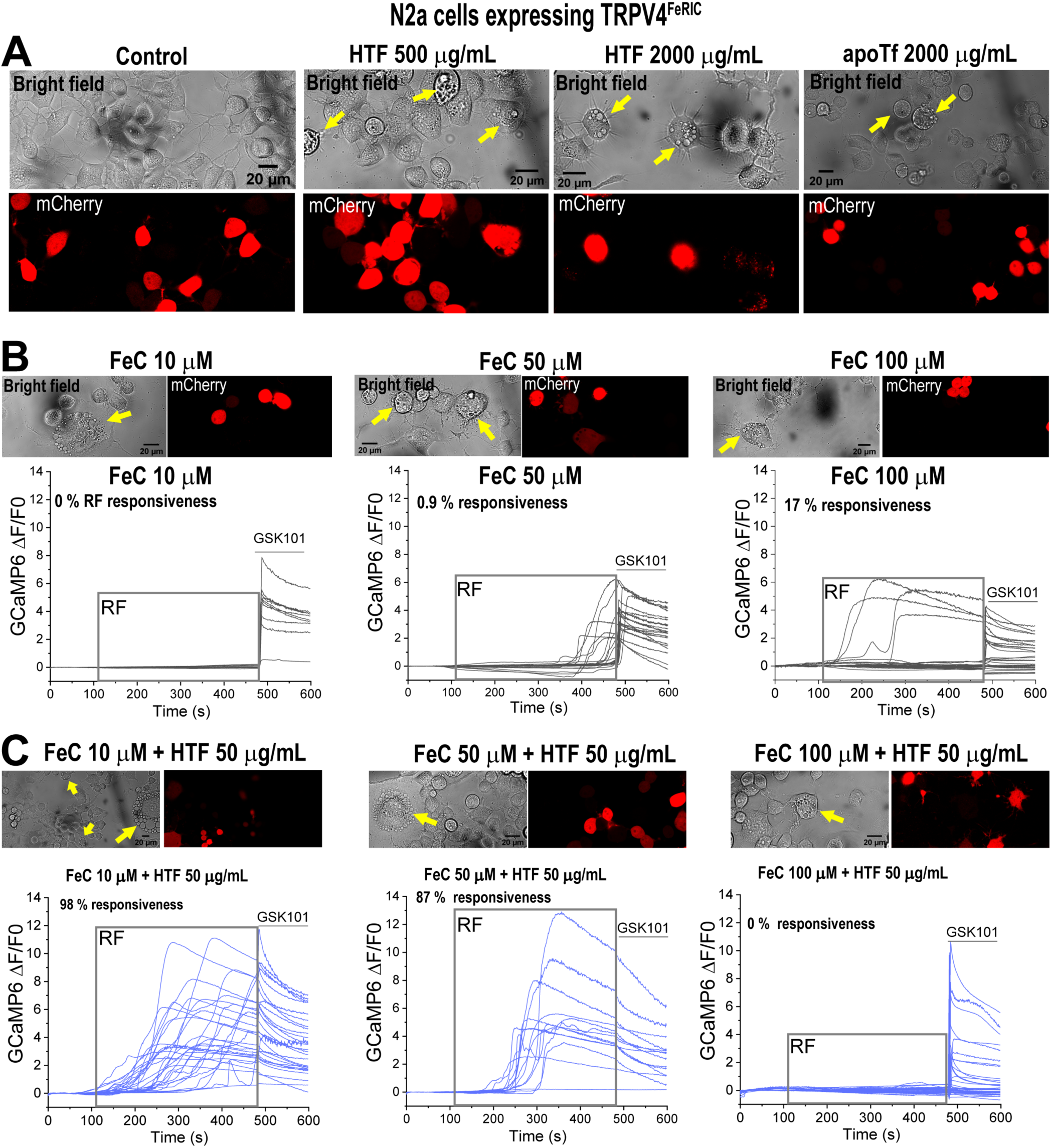
Increasing cellular iron import can increase the RF-induced activation of TRPV4^FeRIC^ but can also negatively affect cell health. Related to Figure 2. (**A**) Bright-field and fluorescence images of N2a cells expressing TRPV4^FeRIC^ (mCherry+). In separate experiments, cells were treated with holotransferrin (HTF, 500 or 2000 µg/mL) or apotransferrin (apoTf, 2000 µg/mL) after transfection. Examples of vacuolated cells are indicated with yellow arrows. (**B, C**) Top images are bright-field and fluorescence images of N2a cells expressing TRPV4^FeRIC^ (mCherry+). Bottom plots are changes in GCaMP6 ΔF/F0 in N2a cells expressing TRPV4^FeRIC^ upon RF stimulation (gray box, 180 MHz, 1.6 μT), followed by GSK101 (solid line). In separate experiments cells were treated with (**B**) ferric citrate (FeC at 10, 50, and 100 μM) or (**C**) a combination of HTF (50 μg/mL) and FeC (10, 50, and 100 μM) after transfection. Examples of vacuolated cells are indicated with yellow arrows. Scale bars = 20 µm.

**Figure S4.**
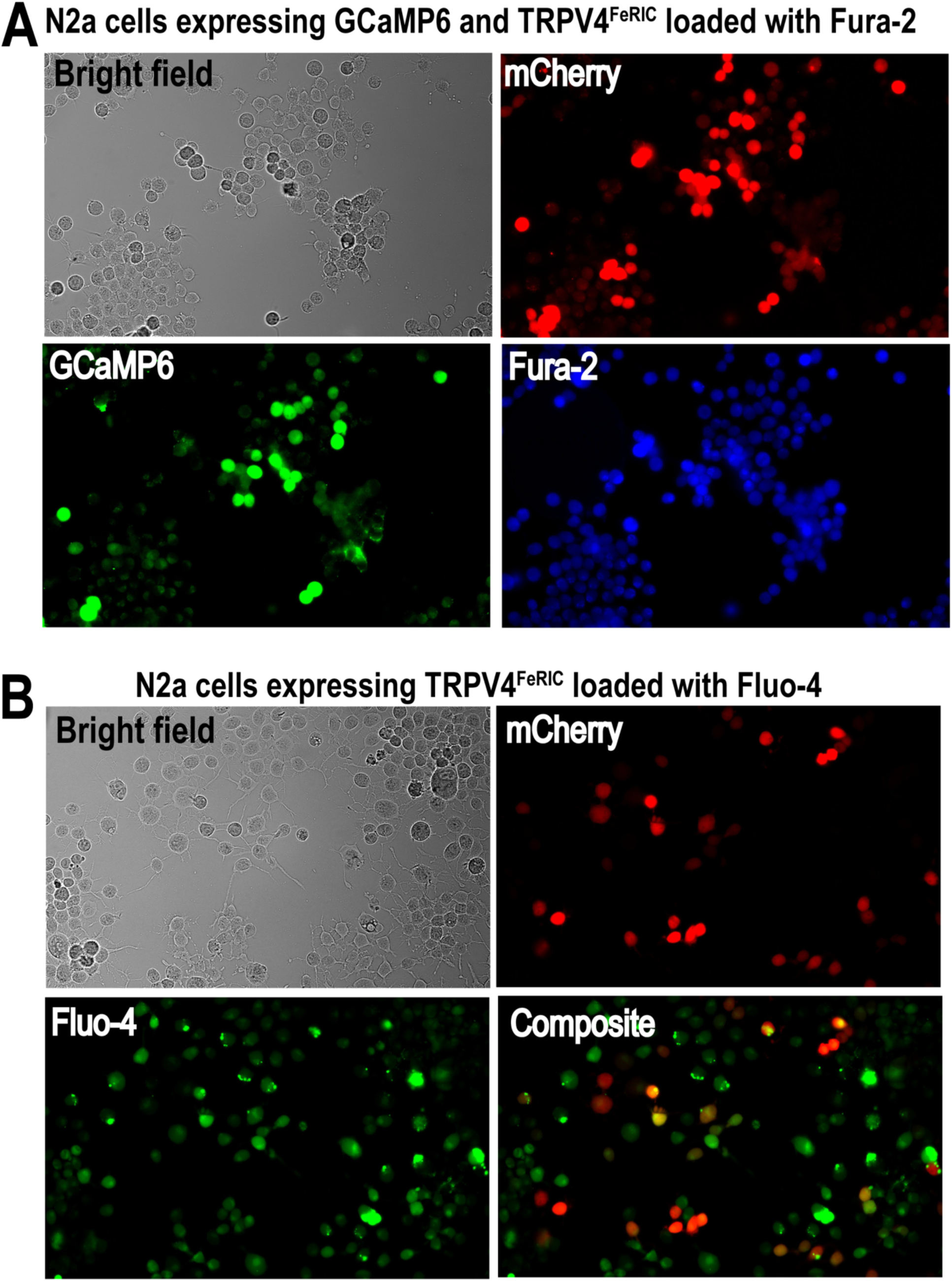
RF-induced activation of TRPV4^FeRIC^ in N2a cells expressing TRPV4^FeRIC^ loaded with Ca^2+^ dyes. Related to Figure 4. (**A**) Representative images of N2a cells expressing GCaMP6 plus TRPV4^FeRIC^ (mCherry^+^) and loaded with Fura-2 (1 μM). (**B**) Representative images of N2a cells expressing TRPV4^FeRIC^ (mCherry^+^) and loaded with Fluo-4 (1 μM).

---

**Table S1.**
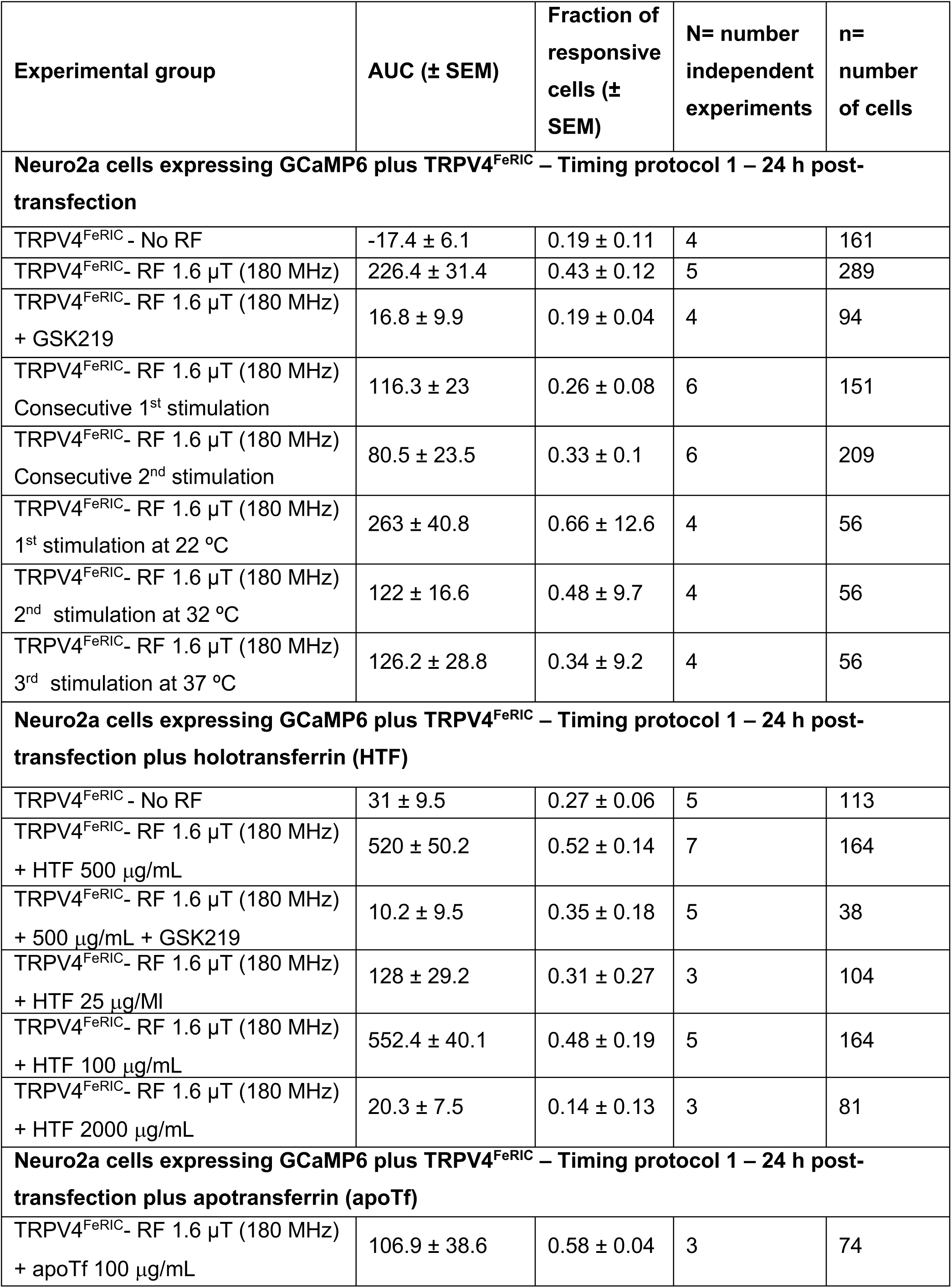
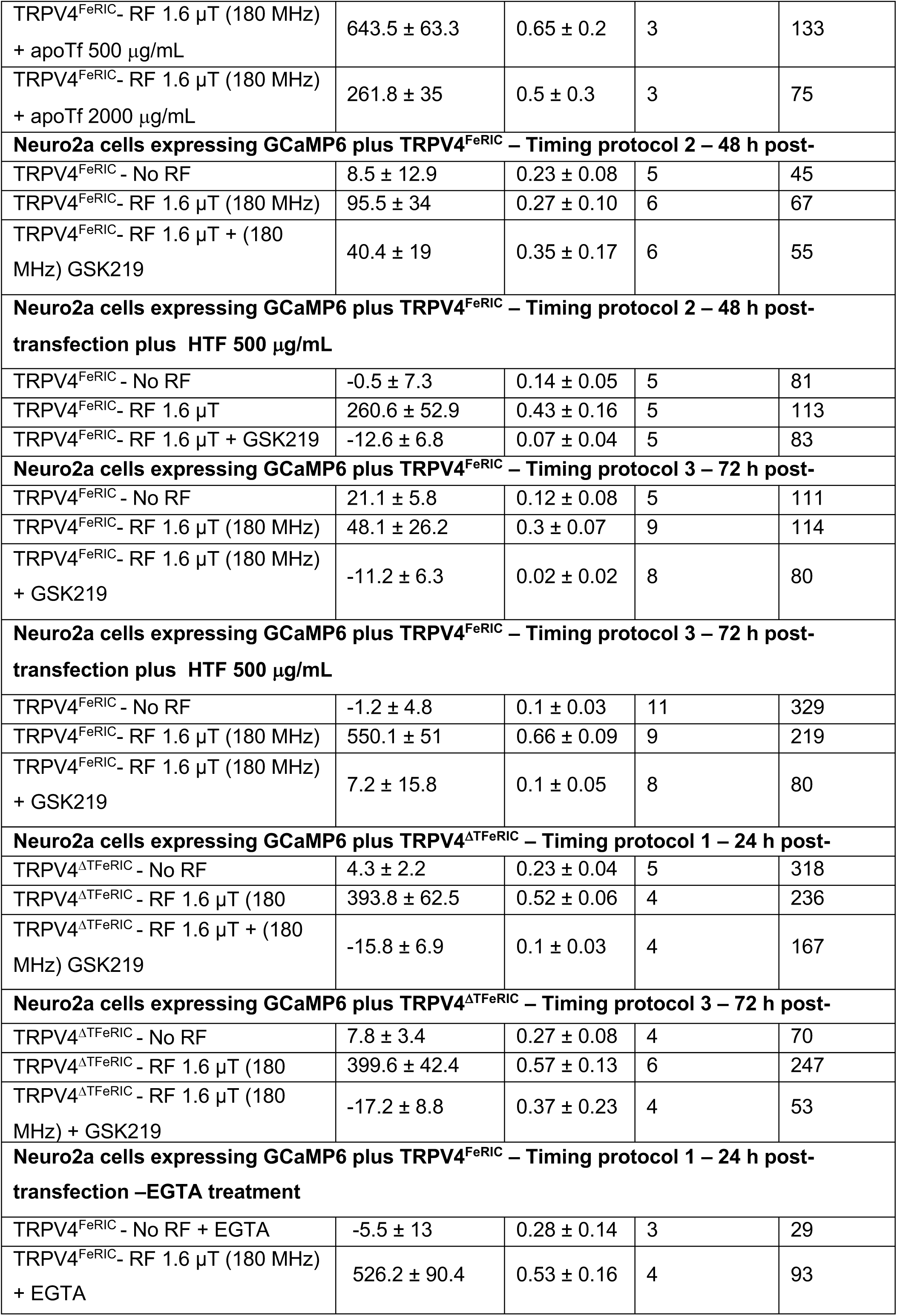
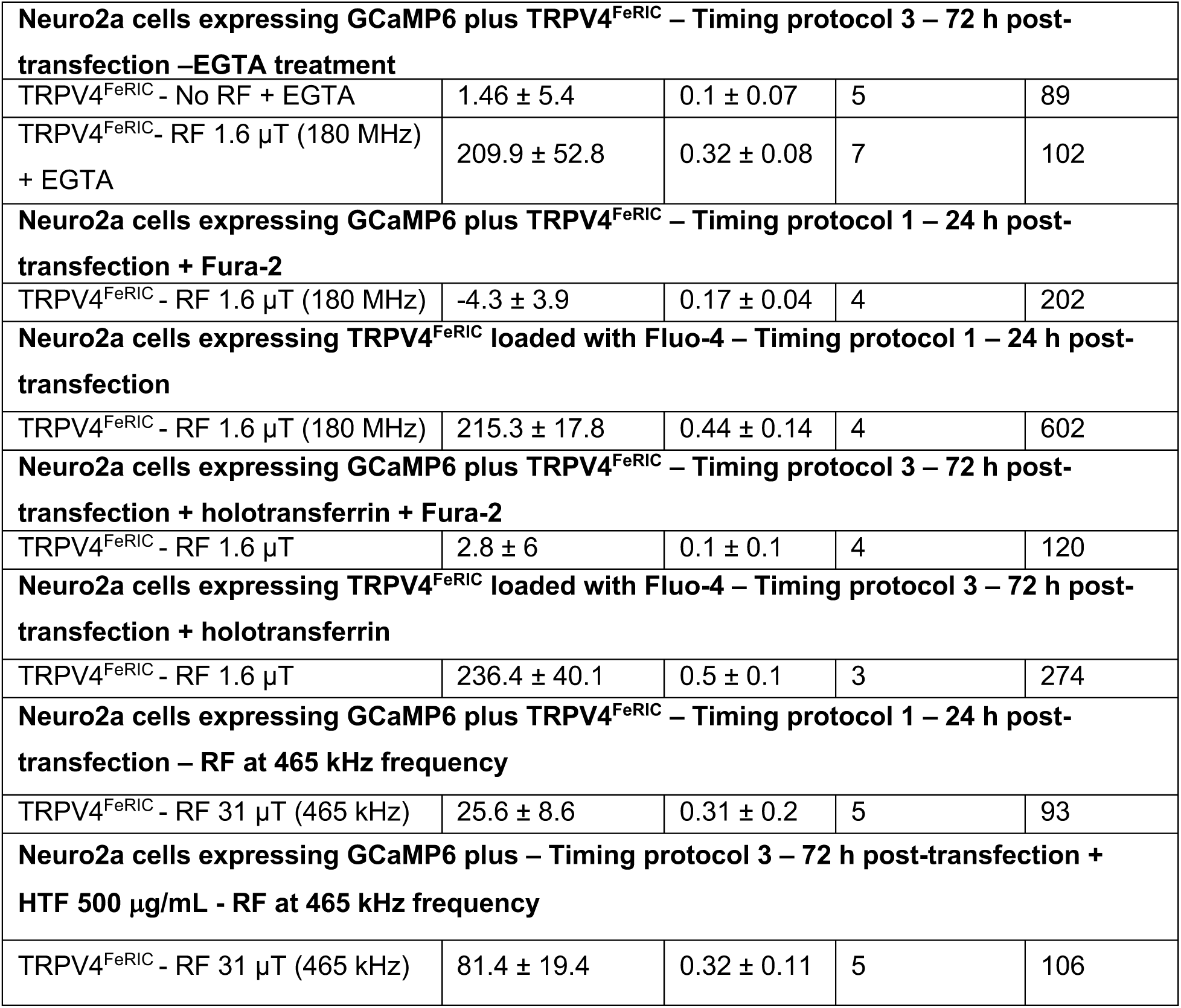
GCaMP6 and Fluo-4 data from N2a cells expressing TRPV4^FeRIC^. Related to Figures 1 - 4. GCaMP6 and Fluo-4 data were quantified as the change in GCaMP6 fluorescence divided by baseline fluorescence (ΔF/F0). For each experimental condition, listed here is the averaged GCaMP6 or Fluo-4 area under the curve (AUC) ± SEM, the fraction of cells responsive to RF, the number of separate experiments (N), and the number of analyzed cells (n).

